# Redox-mediated Kick-Start of Mitochondrial Energy Metabolism drives Resource-efficient Seed Germination

**DOI:** 10.1101/676213

**Authors:** Thomas Nietzel, Jörg Mostertz, Cristina Ruberti, Stephan Wagner, Anna Moseler, Philippe Fuchs, Stefanie J. Müller-Schüssele, Abdelilah Benamar, Gernot Poschet, Michael Büttner, Guillaume Née, Ian Max Møller, Christopher H. Lillig, David Macherel, Iris Finkemeier, Markus Wirtz, Rüdiger Hell, Andreas J. Meyer, Falko Hochgräfe, Markus Schwarzländer

## Abstract

Seeds preserve a far developed plant embryo in a quiescent state. Seed metabolism relies on stored resources and is re-activated to drive germination when the external conditions are favorable. Since the switchover from quiescence to re-activation provides a remarkable case of a cell physiological transition we investigated the earliest events in energy and redox metabolism of *Arabidopsis* seeds at imbibition. By developing fluorescent protein biosensing in intact seeds, we observed ATP accumulation and oxygen uptake within minutes, indicating rapid activation of mitochondrial respiration, which coincided with a sharp transition from an oxidizing to a more reducing thiol redox environment in the mitochondrial matrix. To identify individual operational protein thiol switches, we captured the fast release of metabolic quiescence *in organello* and devised quantitative iodoacetyl tandem mass tag-based (iodoTMT) thiol redox proteomics. The redox state across all Cys-peptides was shifted towards reduction from 27.1 % to 13.0 %. A large number of Cys-peptides (412) were redox-switched, representing central pathways of mitochondrial energy metabolism, including the respiratory chain and each enzymatic step of the tricarboxylic acid cycle (TCA). Active site Cys-peptides of glutathione reductase 2, NADPH-thioredoxin reductase a/b and thioredoxin-o1 showed the strongest responses. Germination of seeds lacking those redox proteins was associated with markedly enhanced respiration and deregulated TCA cycle dynamics suggesting decreased resource efficiency of energy metabolism. Germination in aged seeds was strongly impaired. We identify a global operation of thiol redox switches that is required for optimal usage of energy stores by the mitochondria to drive efficient germination.

## INTRODUCTION

Dry orthodox seeds can preserve their ability to germinate for years, and even millennia in extreme cases (1), which is linked to their minimal metabolic activity when desiccated. At seed imbibition, stored metabolites and enzymes become accessible to reboot metabolic activity in the presence of oxygen. In oilseeds, like those of *Arabidopsis*, the breakdown of triacylglycerol in peroxisomes then progressively provides citrate and succinate as substrate for mitochondrial metabolism (2, 3), which delivers metabolic intermediates, reductant and ATP essential for biosynthesis and cell expansion. Translation of stored transcripts and *de novo* transcription require the successful re-establishment of energy physiology, and only commence in the range of several hours after imbibition (4, 5). This raises the problem that hormonal control relying on genetic programs can only be executed at a stage when physiological, thermodynamic and organizational conditions have already been re-established. Before that, more direct regulatory principles have to apply to first ensure the orderly establishment of cellular metabolism and physiology.

Posttranslational protein modifications have been implicated as a critical regulatory mechanism of metabolic function in the early phases of seed germination and Cys-based redox regulation provides a particularly good candidate mechanism (6, 7). In metabolically reactivated seeds, the reduction of the thiol redox machineries entirely depends on NADPH derived from primary metabolism (8). NADPH provides electrons either to the glutathione/glutaredoxin (GRX)-based redox machineries via glutathione reductases (GRs) or to the different thioredoxins (TRXs) via NADPH-thioredoxin reductases (NTRs) (9).

Already in 1943, reduced glutathione (GSH) could be detected in seed extracts four hours after imbibition, indicating the early establishment of reducing activity, since GSH amounts were much lower in extracts from dry seeds (10). More recently, TRX-dependent reduction of the primary storage proteins in response to imbibition was observed in seeds from different species (11–13), with an impact on the storage protein solubility and accessibility (14). Since TRX-mediated reduction during imbibition was also observed for non-storage proteins of different cellular localization and function (13), the reduction of TRX-linked proteins during seed rehydration is likely a global and conserved phenomenon for orthodox seeds. Specific proteins, such as protein disulfide isomerases and several glutathione S-transferases, were found to be oxidized during imbibition (13), however, which probably reflects the re-establishment of the cell compartment-specific physiological redox environments, i.e. oxidizing in the endoplasmic reticulum. The direct mechanistic connection of Cys redox switching with metabolism raises the hypothesis of Cys redox switching as a fundamental strategy to provide rapid control when hormonal and genetic programmes are not yet fully active. However, testing the plausibility of this hypothesis requires precise information about timing, identity and physiological operation of redox-switched thiol proteins in seeds during metabolic reactivation, which is still missing.

Here, we focus on the earliest events during seed imbibition before hormonal and genetic programmes can dominate germination control. Making use of *in vivo* monitoring of ATP, thiol redox and oxygen dynamics in combination with metabolite analysis, we define the kinetics of re-activation of mitochondrial energy and redox metabolism. Finding that energy and redox re-booting are linked, we dissect the significance of the redox transition in the mitochondria. We capture the transition by establishing a controlled *in organello* model and use iodoTMT-based redox proteomics to identify and quantify individual target cysteines that are operated as thiol switches *in situ*. Using genetic ablation of the proteins most strongly affected we pinpoint the physiological role of the redox transition in maintaining resource efficiency and germination vigor identifying thiol redox regulation as a rapid and direct control strategy.

## RESULTS

### Mitochondrial energy metabolism of seeds starts rapidly at imbibition

To assess the cellular energy dynamics during early germination, we made use of *Arabidopsis* seeds expressing the MgATP^2-^-specific FRET-sensor ATeam 1.03 nD/nA (ATeam) in the cytosol (15). The embryo dominates the sensor signal from the intact seed and the fluorescence intensity was sufficient to be detected through the seed coat **(Fig. 1A)**. Emission spectra showed the characteristic sensor maxima, and the sensor spectrum could be reliably separated from the auto-fluorescence through subtraction of the spectra from seeds without the sensor, both in the dry and the imbibed state **(Fig. 1A, Fig. S1A)**. To monitor cytosolic MgATP^2-^ dynamics live during seed imbibition, intact seeds were rehydrated through online water injection in a multi-well plate reader **(Fig. 1B)**. A rapid increase of FRET directly at imbibition suggested an immediate onset of MgATP^2-^ accumulation in the cytosol **(Fig. 1C)**. The FRET response moved towards a plateau, which was reached at about 70 min (half-maximal sensor response T_1/2_ after 34 ±2 min), suggesting either stable cytosolic MgATP^2-^ concentration or reaching sensor saturation. We observed similar sensor dynamics also by confocal imaging of embryos **(Fig. S1B)**, which could be isolated in an intact state from 10 min after imbibition on. Quantification of adenosine phosphate concentrations in total seed extracts validated a rapid re-establishment of the energy charge (EC) of the adenosine phosphates, which was largely completed after 60 min **(Fig. 1D)**. AMP and ADP levels were dominant in extracts from dry seeds and were decreased by imbibition, while ATP was dominant in imbibed seeds. The EC increased from 0.2 in dry seeds to 0.6 after 1 h imbibition and remained constant afterwards (4 h). As the EC is typically ≥0.9 in fully active cells, the EC is likely to increase further beyond the 4 h time point. A lower apparent EC may alternatively be explained by tissue gradients within the seeds. To assess the source of ATP production and the shift in EC, we next assessed oxygen uptake by the seeds as a measure of mitochondrial respiration. Oxygen consumption was established immediately from the start of the measurement after addition of water, whereas the addition of the respiratory inhibitor cyanide decreased the oxygen consumption **(Fig. 1E)**. We next hypothesized that the fast transition from quiescence to metabolic re-activation may give rise to transient changes of metabolite pools, as a result of fluxes that have not yet reached steady state. Analysis of a set of organic acids associated with mitochondrial energy metabolism **(Fig. 1F)** revealed high malate content in dry seeds, whereas pyruvate, 2-oxoglutarate, succinate, fumarate and lactate were present at lower concentrations. Isocitrate/citrate was below the detection limit, but accumulated to malate levels within 1 h of imbibition and dropped again beyond detection limit at 4 h of imbibition. Succinate and lactate accumulated gradually, possibly reflecting partial hypoxia in central embryo tissues. Pyruvate, 2-oxoglutarate and fumarate pools remained stable. The pronounced isocitrate/citrate transient indicates sequential activity changes within the glyoxylate cycle and the TCA cycle during the first hours of seed imbibition, as mediated by citrate synthase, aconitase, isocitrate dehydrogenase or ATP-citrate lyase. Free amino acid pools were unchanged between dry and imbibed seeds at 1 and 4 h **(Fig. S2)**, suggesting balanced fluxes or no major degradation of storage proteins at this early stage of germination.

**Fig. 1.**
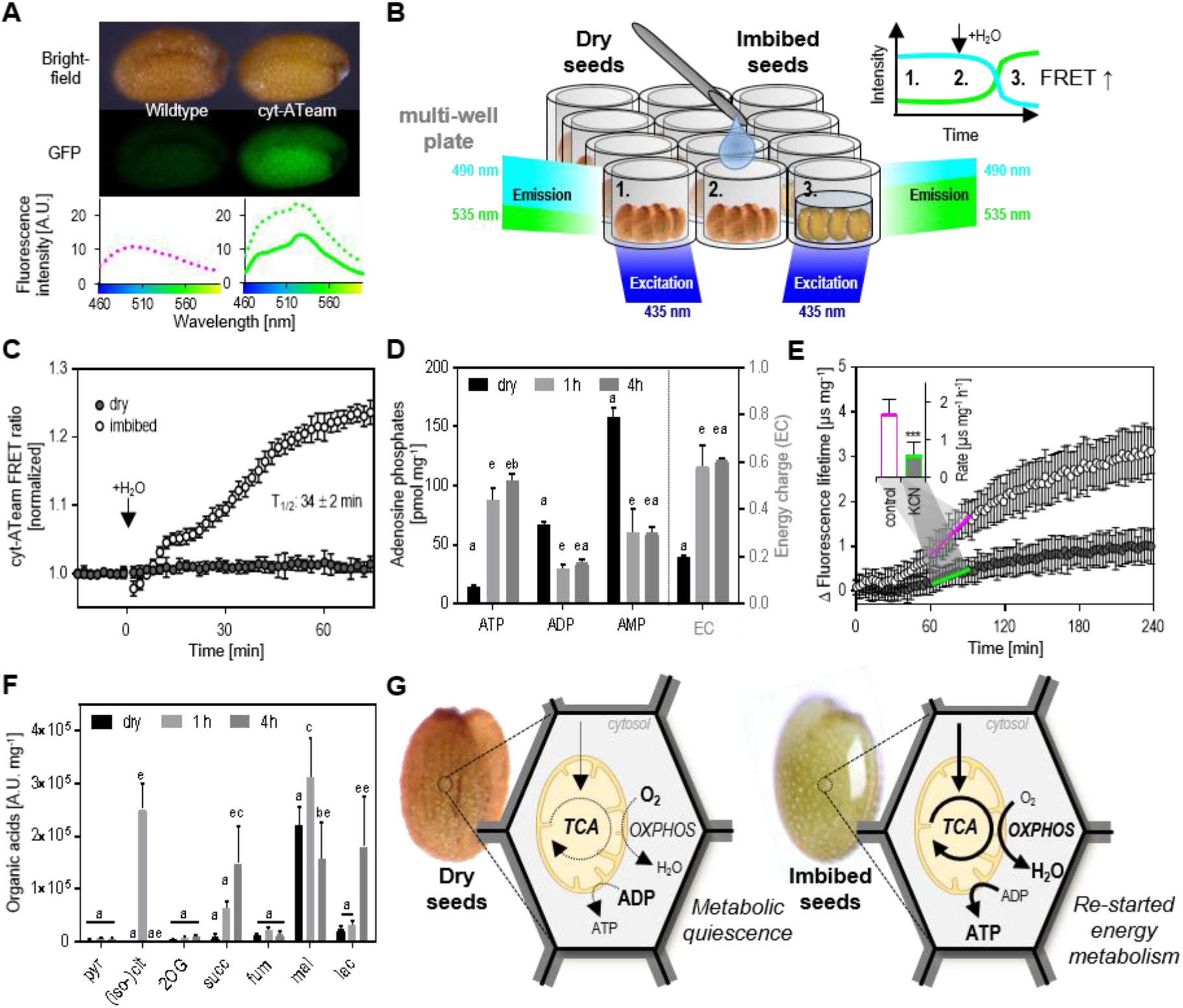
Restart of respiratory energy metabolism in *Arabidopsis* seeds at imbibition. (**A** and **B**) Experimental setup for plate reader-based detection of genetically encoded probes in seeds at imbibition exemplified for the MgATP^2-^ FRET-sensor ATeam. (**A**) Representative brightfield and fluorescence (GFP-filter) images of dry seeds, either with or without expression of cytosolic ATeam (cyt-ATeam). Emission spectra of the seeds are plotted below; dashed magenta line for wildtype (Col-0) seeds, dashed green line for cyt-ATeam seeds and green line for cyt-ATeam seeds corrected for auto-fluorescence of Col-0 seeds (*n* = 3-4 seed batches). (**B** and **C**) Monitoring cytosolic MgATP^2-^ levels in intact seeds through online imbibition and detection of the cyt-ATeam FRET-ratio. 1: dry seed, 2: addition of water, 3. imbibition. The arrows indicate addition of water and T_1/2_ equals the time point of half-maximal sensor response (*n* = 6 seed batches; mean normalized to last value before injection of water ± SD and corrected for Col-0 auto-fluorescence). (**D**) Total ATP, ADP and AMP concentrations in dry seeds and after 1 and 4 h of imbibition (*n* = 4; mean normalized to seed dry weight + SD; two-way ANOVA with Tukey’s multiple comparisons test with a: *p* > 0.05, b: *p* < 0.05 and e: *p* < 0.0001, significant differences between 1 and 4 h imbibition are indicated by a second character). Corresponding energy charge (EC) is ([ATP] + ½ [ADP]) / ([ATP] + [ADP] + [AMP]) is shown on secondary axis (mean + SD). (**E**) Oxygen consumption of imbibed seeds measured with MitoXpress Xtra (fluorescence lifetime of the probe is physically quenched by oxygen). Increase of fluorescence lifetime (Δ FLT) recorded for seeds at control conditions (white circles) or supplemented with 500 μM KCN (grey circles; *n* = 6 seed batches; mean normalized to amount of dry seeds ± SD). Inset shows Δ FLT rate in the timeframe of 60-90 min (two-sided Student’s *t*-test with \*\*\**p* < 0.001) as indicated by the green and magenta line. (**F**) Concentration of organic acids analyzed in dry seeds and after 1 and 4 h of imbibition (*n* = 4; mean normalized to seed dry weight + SD; two-way ANOVA with Tukey’s multiple comparisons test with a: *p* > 0.05, b: *p* < 0.05, c: *p* < 0.01, d: *p* < 0.001 and e: *p* < 0.0001, significant differences between 1 and 4 h imbibition are indicated by a second character). Pyruvate (pyr), (iso-)citrate ((iso-)cit), 2-oxoglutarate (2OG), succinate (succ), fumarate (fum), malate (mal) and lactate (lac). (**G**) Working model of the restart of the respiratory energy metabolism of dry seeds early during imbibition. Width of arrow lines indicate flux rates. Tricarboxylic acid cycle (TCA), oxidative phosphorylation (OXPHOS).

Taken together these data support the idea that mitochondrial respiration is started rapidly with imbibition and before gene expression-based control can underpin hormonal signalling **(Fig. 1G)**. Although the velocity of the activation suggests that the restart is largely driven by physical re-hydration of metabolites and other cell constituents, as well as oxygen supply, early re-booting mitochondrial energy metabolism makes it a likely prerequisite for early metabolic regulation and germination control.

### An early kick-start of the thiol redox machinery

We reasoned that direct regulation through posttranslational protein modifications provides a particularly plausible regulatory framework to ensure orderly progression of early germination before hormonal and transcriptional circuits take control after several hours (16, 17). Focussing on cysteine-linked redox switches, which can be operated rapidly as linked to metabolic electron fluxes, we employed the genetically encoded fluorescent biosensor roGFP2-Grx1 that responds to the glutathione redox potential (*E*_GSH_) (18, 19), to explore the dynamics of thiol redox status specifically in the mitochondrial matrix, i.e. the site of action of most respiratory enzymes **(Fig. 2A)**. Our rationale was that *E*_GSH_ reduction indicates local NADPH provision within the mitochondrial matrix, which is required to reduce GSSG through GR2 in the matrix (20) **(Fig. 2A)**.

**Fig. 2.**
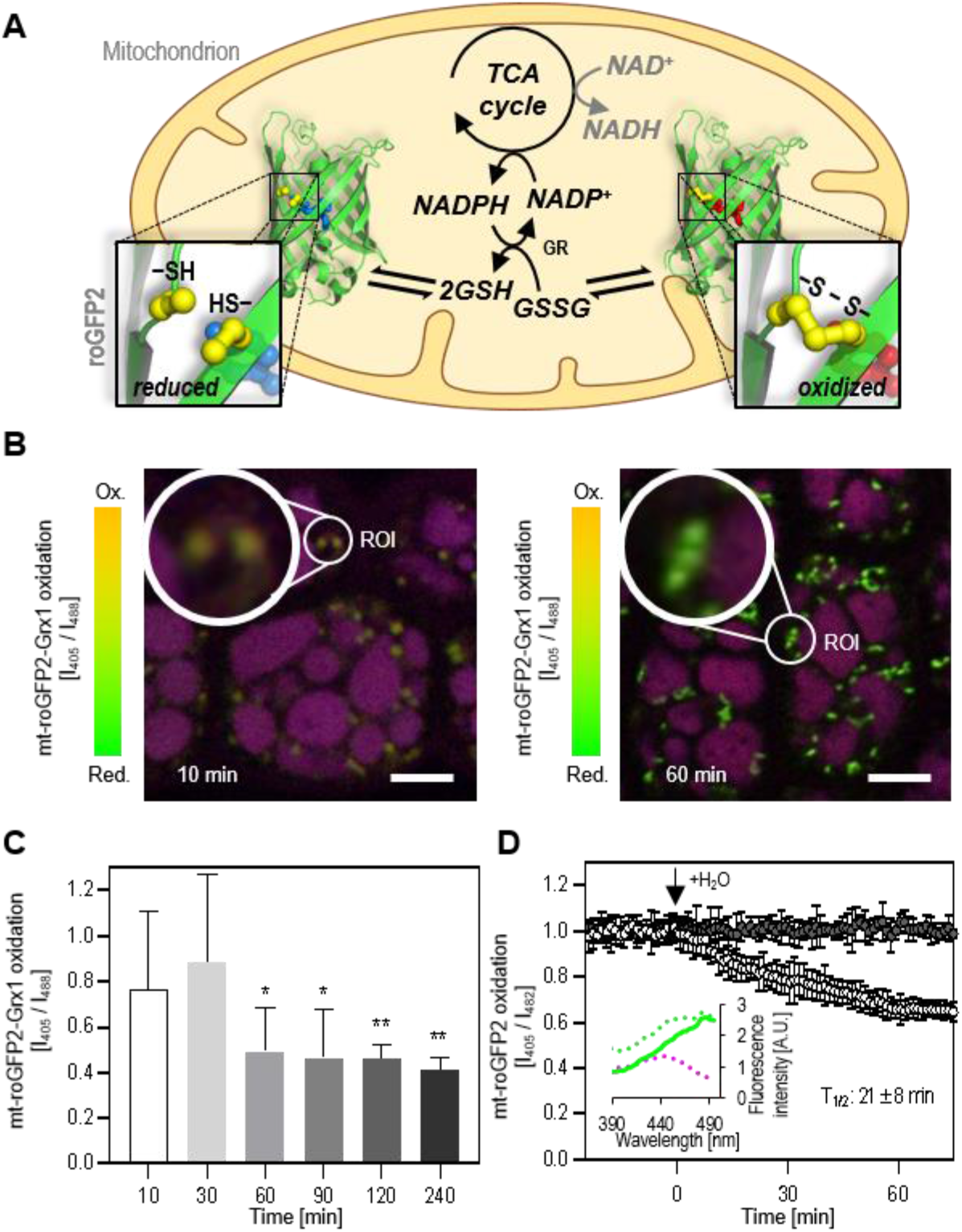
Activation of the mitochondrial thiol redox machinery in *Arabidopsis* seeds at imbibition. (**A**) Reduction of NAD^+^ and NADP^+^ in the mitochondrial matrix is coupled to the activity of metabolic dehydrogenases, including those of the tricarboxylic acid cycle (TCA cycle). NADPH, but not NADH, can act as electron donor to the matrix thiol redox machineries, *e.g*. for the reduction of glutathione disulfide (GSSG) to glutathione (GSH) by glutathione reductase (GR), and to the thioredoxin-system (not depicted). Redox state of the mitochondrial glutathione pool can be monitored by roGFP2 sensors. (**B** and **C**) Change in matrix glutathione redox state in imbibed seeds lacking seed coat and endosperm over time analyzed by confocal microscopy monitoring the 405/488 nm excitation ratio of roGFP2-Grx1. (**B**) Representative confocal images showing embryo epidermal cells imbibed for 10 and 60 min. Regions of interest (ROI) with mitochondria are highlighted and magnified. Overlay of the 405 nm and the 488 nm channels in red and green, protein storage vacuoles appear in purple, due to an auto-fluorescence channel displayed in blue; bars = 10 μm. (**C**) Changes in matrix roGFP2-Grx1 oxidation during seed imbibition analyzed by confocal imaging (*n* = 400-700 individual mitochondria per timepoint in 8-12 individual embryos; mean + SD; two-sided Student’s *t*-test with \**p* < 0.05 and \*\**p* < 0.01). (**D**) Mitochondrial roGFP2 dynamics at imbibition of intact seeds monitored by plate reader-based fluorimetry (see Fig. 1 A and B). The arrow indicates addition of water and T_1/2_ equals the time point of half-maximal sensor response (*n* = 6 seed batches; mean normalized to last value before injection of water ± SD and corrected for Col-0 auto-fluorescence). Inserted panel show excitation spectra of dry seeds, dashed magenta line for Col-0 seeds, dashed green line for mt-roGFP2 seeds and green line for mt-roGFP2 seeds corrected for auto-fluorescence of Col-0 seeds (*n* = 3-4 seed batches).

Similar to imaging of MgATP^2-^, we used ratiometric confocal imaging to measure the dynamics of matrix *E*_GSH_ starting from 10 min after imbibition when intact embryos could be isolated. For separation of the sensor signal from the auto-fluorescence of the protein storage vacuoles, we analyzed several hundred individual mitochondria in several embryos by a region-of-interest analysis (21). Embryos isolated at 10 min showed high 405/488 nm excitation ratios indicating high probe oxidation, which decreased markedly after 60 min, indicating a pronounced transition towards probe reduction **(Fig. 2B,C)**. To control for potential effects of embryo isolation, we adjusted the plate reader-based analysis protocol **(Fig. 1B)** for roGFP2 to assess the probe response in intact seeds. Despite a low signal-to-noise ratio due to a high proportion of auto-fluorescence in the mitochondrial sensor line, the characteristic excitation spectrum of roGFP2 was reliably recorded **(Fig. 2D, Fig. S3A)** to monitor the redox dynamics in mitochondria of intact seeds. A rapid sensor reduction was detectable at imbibition within the first minutes that reached a plateau within 60 min (half-maximal sensor response T_1/2_ after 21±8 min), indicating rapid establishment of a highly reduced glutathione pool **(Fig. 2D)** as typically found in the mitochondria of vegetative plant tissues (19). Rapid *E*_GSH_ reduction was also observed using a cytosolic sensor, although with distinct, biphasic kinetics **(Fig. S3B)**. The sensor reduction dynamics in both subcellular compartments coherently indicate that the thiol redox machineries are restarted very early at imbibition.

Rapid and simultaneous re-establishment of subcellular energy and redox physiology through mitochondrial activity may be a characteristic of germinating, orthodox seeds **(Fig. 1, 2)**, and a prerequisite to drive and regulate subsequent steps of germination. Specifically, activation of the thiol redox machinery in the mitochondrial matrix provides a mechanism to reset the landscape of operational protein thiol switches to control mitochondrial function.

### Re-booting the thiol redox machinery in purified mitochondria

To reveal which specific protein thiol switches are operated within the functional mitochondria we developed a strategy based on quantitative redox proteomics. Maintaining meaningful thermodynamic, kinetic and spatial constraints for the operation of thiol switches was a key priority, since kinetic control delivers specificity in thiol redox regulation (22), and cannot be straightforwardly studied *in vitro* (23). Since whole seed extracts are dominated by storage proteins, leaving mitochondrial proteins strongly underrepresented in proteome analyses (24), we optimized a model system to capture the fast release of metabolic quiescence that we observed in seeds using purified seedling mitochondria. Respiratory metabolism is quiescent in dry seeds due to lack of free water for substrate solubilization and enzyme activity; consequently, no NADPH is generated in the matrix and thiols cannot be effectively maintained in a reduced state, as indicated by the oxidized roGFP2 sensor **(Fig. 2)**. Analogously, isolated plant mitochondria are metabolically quiescent due to respiratory substrate depletion and cannot generate NADPH **(Fig. 3A)**.

**Fig. 3.**
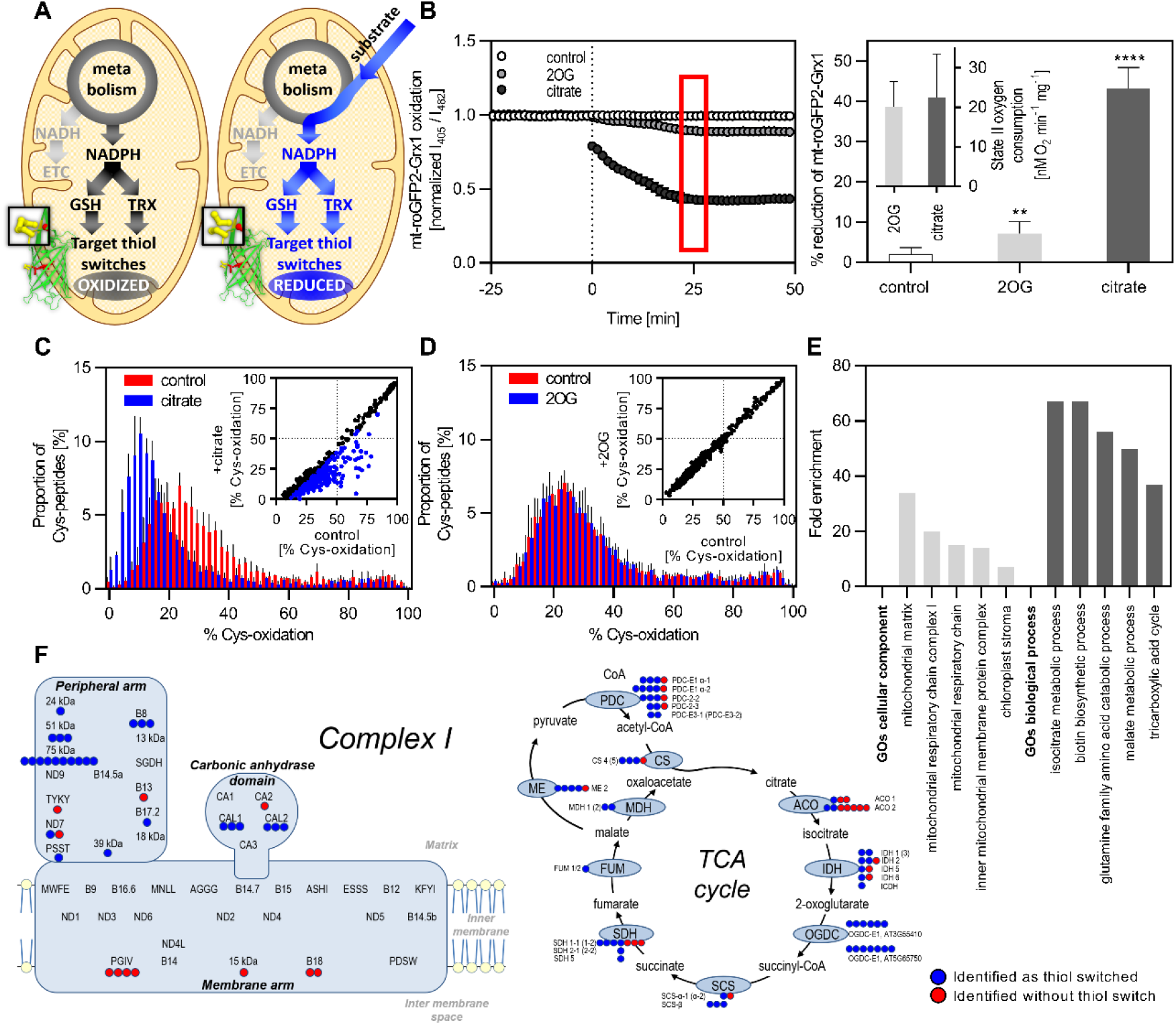
Re-booting the thiol redox machinery of isolated mitochondria to define the proteomic landscape of mitochondrial proteinaceous Cys redox switches. (**A**) Schematic model of quiescent mitochondria and their metabolic re-activation by substrate addition. Substrates are metabolized reducing NAD^+^ and NADP^+^. Both NADH and NADPH can be oxidized by the electron transport chain (ETC), but only NADPH can act as reductant for the matrix thiol redox machineries. Matrix-targeted roGFP2-Grx1 acts as artificial target of the glutathione redox machinery. (**B**) Representative roGFP2-Grx1 reduction kinetics of quiescent mitochondria, either supplemented with 10 mM 2-oxoglutarate (2OG) or 10 mM citrate (*n* = 4 technical replicates, mean ± SD; dashed line indicates addition of substrates and red box highlights the oxidation state after 25 min). The mt-roGFP2-Grx1 redox state calculated for each metabolic condition after 25 min in the right panel; for calibration of the sensor, mitochondria were incubated with 5 mM dipyridyldisulfide (DPS, oxidant) or 20 mM dithiothreitol (DTT, reductant) for 20 min (twosided Student’s *t*-test with \*\**p* < 0.01 and \*\*\*\**p* < 0.0001). Inset shows state II oxygen consumption of isolated mitochondria supplemented either with 2OG or citrate (*n* = 5, mean + SD). (**C** and **D**) Abundance profiles of the redox states of Cys-peptides in quiescent and respiring mitochondria. Isolated mitochondria supplemented with citrate (**C**) or 2OG (**D**) and incubated for 25 min before differential thiol labeling with iodoTMTs. Redox states of individual peptides quantified for isolated mitochondria with and without respiratory substrate (blue and red bars, respectively). The distribution of cysteine peptide oxidation levels is shown by the proportion of the total number of peptides in each 2 % quantile of percentage oxidation (*n* = 3, mean + SD). In insets, percentage of Cys-oxidation without substrate addition is plotted against percentage of Cys-oxidation after substrate addition for individual Cys-peptides. Cys-peptides with significant change in their Cys-oxidation state are displayed in blue (*t*-test corrected for multiple comparisons by Benjamini, Krieger & Yekutieli with < 2 % FDR). (**E**) The five most enriched GO terms of the ‘*cellular component*’ and ‘*biological process*’ categories of a PANTHER-overrepresentation analysis of all significantly redox-switched Cys-peptides. (**F**) All identified Cys-peptides of the individual subunits of the mitochondrial Complex I mapped on the schematic structure according to Braun *et al*., 2014 (left). All Cys-peptides identified for enzymes of the TCA cycle and closely associated (right). Blue: significantly redox-switched Cys-peptide; Red: not significantly redox-switched Cys-peptide.

To recreate the rapid redox transition as observed in mitochondria of intact seeds we supplied purified mitochondria, expressing mitochondrial roGFP2-Grx1, with metabolic substrates. Different matrix dehydrogenases couple substrate oxidation to the reduction of NAD^+^ and NADP^+^ (25). Yet, only NADPH can act as an electron donor to the thiol redox machinery **(Fig. 3A)**. Citrate feeding triggered a pronounced reduction of the sensor **(Fig. 3B, left panel)**. After 25 min, the sensor signal reached a plateau at a highly reduced state. The rapid and pronounced sensor response indicates active and sustained NADPH generation in the matrix, with the NADP-dependent isoform of isocitrate dehydrogenase (ICDH) as likely source. As a control substrate, we used 2-oxoglutarate (2OG), for which there is no obvious NADPH-producing enzyme downstream (25). 2OG triggered only a weak reduction of the sensor. Mitochondrial oxygen consumption rates triggered by the two substrates were comparable, indicating that both substrates were taken up into the matrix and metabolized at similar rates **(Fig. 3B, inset in right panel)**. Taken together the data suggest that 2OG oxidation predominantly produced NADH, while citrate oxidation caused significant rates of NADPH production in addition. The data further suggest that the redox states of NAD and NADP pools are separated in the matrix, consistent with the lack of any mitochondrial transhydrogenase protein homologue in higher plant genomes.

### The landscape of operational cysteinyl thiol redox switches of the mitochondrial matrix

Next, we aimed to identify those mitochondrial thiol switches that are operated *in situ* during the transition from quiescence to activity of respiratory matrix metabolism. We developed and optimized fast differential Cys labeling in intact mitochondria using thiol-specific iodoTMTs. Differential iodoTMT-based redox proteomics allows to quantify the degree of oxidation (% oxidation) of individual proteinaceous cysteinyl thiols by tandem mass spectrometry (MS/MS). Absolute quantitation of the Cys redox landscape at Cys-peptide resolution was not explored by previous redox proteomic approaches in plants. Previous approaches typically pinpointed redox changes at the protein level and/or rather focussed on the relative percentage of thiol redox switching (Δ % oxidation) as induced by a treatment. Δ % oxidation can also be determined as a difference between the absolute % oxidation of the two metabolic states. The iodoTMT approach as applied here does not resolve the chemical identity of Cys modification, however, which will include several different types of oxidative Cys-peptide modifications, such as disulfides, sulfenylation and nitrosylation. Potential changes in protein abundance between individual replicates are accounted for by internal normalization of the differential labeling approach. Protein abundance remained stable **(Fig. S4)**. The comparison between quiescent and citrate-respiring mitochondria (sampled after 25 min) allowed for the quantitation of 741 Cys-peptides, mapping to 425 different proteins (734 Cys-peptides mapping to 443 proteins for the control using 2OG supplementation) **(Dataset S1)**. Activating mitochondrial metabolism with citrate resulted in a global reductive shift from 27.1 % Cys-oxidation on average under quiescence to only 13.0 % Cys-oxidation (Δ14.1 %; **Fig. 3C**). By contrast, inducing respiration with 2OG did not change Cys-oxidation significantly (27.1 % under quiescence to 27.0 % with 2OG; Δ0.1 %) and the characteristic distribution of the redox states of Cys-peptides was fully preserved **(Fig. 3D)**. The individual Cys-peptides responded highly differentially to citrate respiration, with 412 peptides mapping to 245 different proteins undergoing a significant reductive shift, while no peptide underwent oxidation **(Fig. 3C inset; Fig. S5D)**. By contrast not a single Cys-peptide was significantly shifted in its degree of oxidation in response to 2OG respiration **(Fig. 3D inset; Fig. S6D)**. Those effects were highly reproducible as evidenced by very similar patterns across the independent replicates **(Fig. S5A,B; Fig. S6A,B)**. The pronounced reductive shift that we had observed fluorimetrically under citrate respiration using matrix roGFP2-Grx1 **(Fig. 3B)** was also observed in the proteomic approach, where a roGFP2 Cys-peptide was detected and showed a reductive shift of Δ30.4 % **(Dataset S1)**.

The individual thiol-switched Cys-peptides differed strongly in their degree of redox shifting **(Dataset S1)**. The Cys-peptides showing the most pronounced reductive shifts mapped to the plastidial/mitochondrial GR2 (Δ61.7 %) (20), the cytosolic/mitochondrial NADPH-dependent thioredoxin reductases a and b (NTR a/b; Δ49.1 %) (26, 27) and mitochondrial TRX-o1 (Δ48.1 %) (28). All three peptides included the catalytic cysteines of their proteins, mirroring their role as *active* thiol redox enzymes, consistent with previous findings of particularly reactive catalytic cysteines (29, 30). Since two Cys-residues were present in all three peptides, the quantified redox shift represents an average and may be even larger for one of the two cysteinyl thiols. However, over 80 % of Cys-peptides contained a single Cys making the quantification unambiguous for that specific site **(Fig. S7A)**.

To investigate if, in addition to the thiol redox machineries, specific biological functions were preferentially associated with those thiol switches that were operational *in situ*, we used a gene ontology (GO) enrichment analysis (http://geneontology.org). Despite bias through the input of enriched mitochondrial fractions, the analysis has proven useful to highlight overrepresented pathways (30). Central components and processes of the mitochondrial respiratory metabolism were enriched, including those of the mitochondrial respiratory chain and the TCA cycle **(Fig. 3E)**, suggesting central respiratory metabolism harbours hotspots of operational thiol switching. The enrichment of ‘chloroplast stroma’ proteins can be accounted for by a combination of plastidic contamination **(Fig. S7B)** and by proteins assigned as stromal that are actually also mitochondrial. An example is GR2, which is annotated as plastidic, but was demonstrated to be dual-targeted, also to the mitochondria (20). Further functions represented by several different proteins with active thiol redox switches included iron-sulfur metabolism, amino acid metabolism and mitochondrial RNA processing (PPR proteins) (**Dataset S1**).

As an independent way to identify hotspots of operational thiol switching in the intact mitochondrion we looked for proteins for which more than one thiol-switched Cys-peptide was identified. 94 proteins contained two or more Cys-peptides that were significantly redox-switched. The 75 kDa subunit (AT5G37510) of the peripheral arm of Complex I contained even ten functional thiol redox switches **(Fig. 3F)**. Since Complex I was also pinpointed by the GO term analysis **(Fig. 3E)**, we further assessed the locations of the redox-switched Cys within the complex. The relevant proteins were all part of the peripheral arm (21 Cys-peptides switched, 3 non-switched) and the plant-specific carbonic anhydrase domain (6 Cys-peptides switched, 1 non-switched), which may be partly explained by accessibility to soluble redox enzymes. In contrast, all seven identified Cys-peptides of the membrane arm were exposed to the intermembrane space (IMS) (31) and were not redox-switched, which validates the experimental system in which citrate feeding provides NADPH in the matrix, but not the IMS.

Recent work demonstrated Trx-mediated redox regulation of two TCA cycle enzymes, succinate dehydrogenase and fumarase (28). Our dataset validates the presence of operational thiol switches on both players and pinpoints operational Cys switches. It further expands the current picture by the identification of at least one active thiol switch for each enzymatic step of the TCA cycle **(Fig. 3F)**, as a mechanistic basis for comprehensive redox control of this hub of central metabolism. The E1-subunit of the oxoglutarate dehydrogenase complex (OGDC) was identified with seven redox-switched Cys-peptides (AT5G65750, three non-unique peptides also map to the isoform AT3G55410). Fourteen Cys-peptides from different pyruvate dehydrogenase complex (PDC) subunits were redox-shifted. For the succinate dehydrogenase complex (SDH) six out of nine identified Cys-peptides were redox-switched, while only three out of eight Cys-peptides identified from aconitase (ACO) 1 and 2 were redox-switched.

The proteomic compendium of active mitochondrial thiol switches that we have generated demonstrates that a large number of proteinaceous Cys-thiols can not only be reduced *in vitro* (32–36), but are operated in the isolated mitochondrion linked to metabolic activity. All shifts were reductive, reflecting the influx of electrons into the matrix thiol redox systems, and quantitative, since most peptides were maintained partially reduced in the quiescent metabolic state. It is likely that the operation of a thiol switch *in situ* is typically not an all-or-nothing process, although even a partial redox shift of an individual Cys on mammalian Uncoupling Protein 1 (Cys253) by less than 10% has been shown to be able to trigger dramatic metabolic changes (37). Yet, more conceivably in seed imbibition the concerted operation of many switches on many different enzymes at once may have a considerable impact on the fluxes through mitochondrial energy metabolism.

### Mutants of the mitochondrial thiol redox machinery are impaired in germination

To assess the physiological significance of the rapid reduction of the mitochondrial thiol redox machinery during early germination, we aimed to understand its impact on seed energy metabolism and germination characteristics. Since the most strongly thiol-switched peptides belonged to the catalytic sites of GR2, NTR a/b and TRX-o1, we selected *Arabidopsis* mutant lines lacking those proteins (20, 27, 28). In addition, those proteins operate upstream in potential redox regulation cascades, enabling an integrated picture of thiol switching on several target enzymes at once, even though a degree of compensation between the thiol redox systems needs to be anticipated, based on previous observations (20, 27, 38).

Strikingly, oxygen consumption rates early after imbibition (here measured between 60 to 90 min) were strongly increased in the seeds of all three mutants **(Fig. 4A)**, indicating increased respiration rates with the potential of boosting oxidative phosphorylation and ATP synthesis rates. Yet, there was no consistent impact on the different adenosine phosphate pools. Only *ntr a/b* showed a change in EC after 1 h of imbibition, which was even lowered, however. The adenylate pools of the other seeds recovered as in Col-0 **(Fig. 4B)**. Increased respiration rates coinciding with unchanged adenylate charge clearly point to less efficient ATP generation or higher consumption. To test the hypothesis of de-regulated mitochondrial enzymatic activity as a common cause of inefficient respiration in the mutants, we assessed the free organic acid pools of the seeds of all lines.

**Fig. 4.**
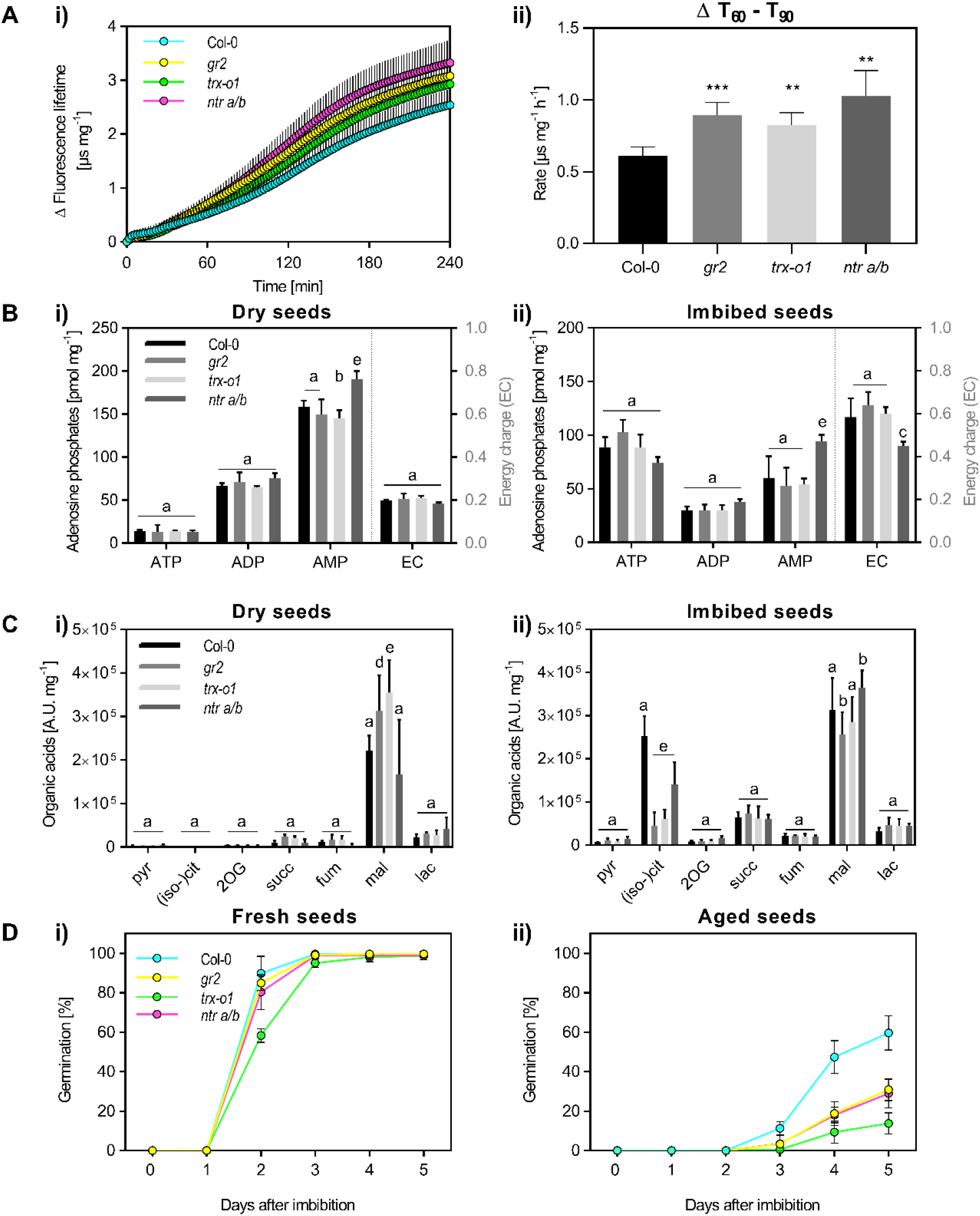
The effect of impairing the mitochondrial thiol redox machinery on energy metabolism and germination. Synchronized seeds of Col-0, *gr2*, *trx-o1* and *ntr a/b* lines were analyzed for their performance during germination. (**A**) Oxygen consumption of imbibed seeds measured with MitoXpress Xtra. **i**) oxygen uptake as indicated by increase in fluorescence lifetime (Δ FLT) over time (*n* = 5 seed batches; mean normalized to amount of dry seeds + SD). **ii**) Δ FLT rate in the 60-90 min time window (two-sided Student’s *t*-test with \**p* < 0.05, \*\**p* < 0.01 and \*\*\**p* < 0.001 if compared with Col-0). (**B**) Total ATP, ADP and AMP concentration (**i**) in dry seeds and (**ii**) after 1 h of imbibition (*n* = 4; mean normalized to seed dry weight + SD; two-way ANOVA with Dunnett’s multiple comparisons test with b: *p* < 0.05, c: *p* < 0.01, d: *p* < 0.001, e: *p* < 0.0001 if compared with Col-0). Corresponding energy charge (EC) is ([ATP] + ½ [ADP]) / ([ATP] + [ADP] + [AMP]) is shown on secondary axis (mean + SD). (**C**) Concentration of organic acids analyzed (**i**) in dry seeds and (**ii**) after 1 h of imbibition (*n* = 4; mean normalized to seed dry weight + SD; two-way ANOVA with Dunnett’s multiple comparisons test with b: *p* < 0.05, c: *p* < 0.01, d: *p* < 0.001 and e: *p* < 0.0001 if compared with Col-0). Pyruvate (pyr), (iso-)citrate ((iso-)cit), 2-oxoglutarate (2OG), succinate (succ), fumarate (fum), malate (mal) and lactate (lac). (**D**) Germination over 5 days (**i**) for freshly harvested seeds and (**ii**) after a controlled deterioration treatment (CDT) by storage at 37 °C and 75 % rel. humidity for 19 days to mimic seed aging. The statistical analysis of germination rates of *gr2*, *trx-o1* and *ntr a/b* as compared to Col-0 seeds is shown in **Table S1**.

In dry seeds the different pools were unchanged; only malate content differed between the lines, albeit without a consistent pattern **(Fig. 4Ci)**, indicating a similar metabolic situation at the start of imbibition. As observed in Col-0 **(Fig. 1F)**, the organic acid pools responded markedly to imbibition, and strikingly a consistently lowered (iso-)citrate content was observed for all mutants as compared to Col-0 at the 1 h time point **(Fig. 4Cii)**. Since (iso-)citrate accumulation occurs as a transient **(Fig. 1F)**, the modification of its amplitude and/or timing shows that the flux into and/or out of the (iso-)citrate pool is affected in the mutants and provides evidence for de-regulated respiratory carbon metabolism in the thiol redox mutants during early germination. Of the 19 free amino acid pools, 16 did not differ between the lines and remained stable at imbibition **(Fig. S8)**. As an exception, Glu, Asn and Pro were slightly increased in dry *ntr a/b* seeds and accumulated in *gr2* seeds at imbibition, indicating that de-regulation of amino acid metabolism was limited to specific pathways, some of which are located in the mitochondria (39).

To investigate the potential impact of respiratory de-regulation on seed germination, we analyzed germination vigor in freshly harvested seeds over five days after imbibition. All lines showed efficient germination with 100 % germination rate within five days. Only *trx-o1* showed temporarily lowered germination efficiency after two days **(Fig. 4Di, Table S1)**. The potential defect in *trx-o1* was consistent with a previous observation (28), where seeds showed much lower germination vigor overall, however. We hypothesized that, in fresh seeds, efficient carbon metabolism may not be limiting and investigated the germination characteristics of seeds that were exposed to a controlled deterioration treatment (CDT). CDT at 37 °C, 75 % rel. humidity for 19 days decreased germination vigor for all lines, and only 60 % of Col-0 seeds germinated within five days **(Fig. 4Dii)**. Yet, all three redox mutants showed much stronger defects in germination ability and this effect was observed consistently across different aging regimes **(Fig. S9; Table S2)**. Taken together the different results demonstrate the importance of mitochondrial thiol redox switching in the efficient use of stored resources by respiratory metabolism in germinating seeds.

## DISCUSSION

Aiming to understand mechanisms that underpin the sharp metabolic transition from seed quiescence to the activation of germination, our fluorescent sensing data demonstrate that MgATP^2-^ synthesis and oxygen consumption start almost instantaneously upon rehydration. They illustrate how rapidly the transition from quiescence to metabolic activity occurs and highlight the involvement of mitochondrial respiration **(Fig. 1C-F)**, which requires structurally intact and functional mitochondria in the desiccated embryo cells. This, as well as our observation of fluorescently labeled mitochondria in embryo cells after 10 min of imbibition **(Fig. 2B)**, is in contrast to the previous postulate that a maturation phase is required for respiratory incompetent proto-mitochondria before respiration can commence (40), but in agreement with recent observations in pea and *Arabidopsis* seeds (41, 42). Work in lettuce seed extracts has pinpointed the early establishment of adenylate charge, but the *in vivo* dynamics of the response could not be resolved (43). Oxidative phosphorylation sets the balance between the adenylate pools together with adenylate kinase (41), explaining the rapid conversion of AMP to ATP **(Fig. 1D, 4B)**. While oxidative phosphorylation appears to dominate within the first hour, the accumulation of lactate between 1 and 4 h suggests a contribution by fermentation to ATP synthesis **(Fig. 1F)**. That makes the rapid establishment of respiratory energy homeostasis a likely prerequisite for other early germination events, such as translation or *de novo* transcription, which underpin major hormonal checkpoints.

RoGFP-based *in vivo* sensing suggests that re-establishment of the thiol redox status of the mitochondrial matrix and the cytosol is intimately linked to the re-establishment of energy metabolism **(Figs. 1, 2)**. Low EC coincides with oxidized glutathione status, which reflects the absence of sufficient metabolic flux to provide phosphorylation potential and reductant in the form of NADPH. The rapid re-activation of the thiol redox machinery is consistent with the observation of strongly increased GSH levels in seed extracts 4 h after imbibition (10). Further, our data demonstrate that an active thiol redox machinery is not only a side effect of metabolic re-activation, but plays an important regulatory role in germination **(Fig. 4)**. Mutants of three different genes (*gr2, ntr a/b, trx-o1*), all of which are part of the mitochondrial thiol redox machinery, consistently showed increased respiratory rates in the presence of unaltered adenylate dynamics, suggestive of lower efficiency of energy conservation from respiring stored carbon resources. Specific de-regulation of (iso-)citrate is consistent with this interpretation since major citrate and succinate flux is expected from peroxisomal triacylglycerol degradation in *Arabidopsis* seeds (2). Based on our data we propose a model, in which the dampened (iso-)citrate transient at the 1 h time point is caused by de-repressed efflux from the (iso-)citrate pool **(Figs. 1F, 4C)**, leading to increased respiration rates **(Fig. 4A)** and carbon loss in the mutants. While fresh seeds can tolerate inefficient mitochondrial metabolism up to a degree, aging further compromises seed mitochondria (44) leading to drastically decreased germination vigor **(Fig. 4D)**. Our observation that every enzymatic step of the TCA cycle features at least one operational thiol switch **(Fig. 3F)** makes it likely that several TCA cycle proteins contribute to the overall activity pattern of the pathway. Interestingly, a repressive effect by TRX-o1-mediated reduction on activity has recently been found for the TCA cycle enzymes succinate dehydrogenase and fumarase (28). For two mutants of the TRX-based redox machinery similar germination defects were observed. Our model provides a plausible explanation for the data, but it is unlikely to be comprehensive, due to the central role of mitochondrial respiration and carbon metabolism in the cell, which means that defects likely trigger further pleiotropic rearrangements. The start from a joint quiescent situation (dry seeds), may minimize such effects, as supported by the measured metabolite profiles **(Fig. 4)**, but differences already set during seed filling and/or maturation cannot be ruled out. For instance, a wrinkled seed phenotype was reported for the *ntr a/b* mutant, which was shown to be due to role of NTR a/b in the cytosol and not in the mitochondria, however (27).

Several candidate targets for the plant mitochondrial TRX and GRX machineries have been identified by *in vitro* trapping approaches using protein extracts (32–34, 36). Those approaches have been instrumental in identifying thiol proteins that can react with TRX and GRX active site thiols in principle. However, they cannot account for micro-compartmentalization, kinetic competition and thermodynamic conditions, which are key constraints governing thiol redox regulation *in vivo* (23). *In situ* trapping approaches, as performed in mitochondria of human embryonic kidney cells (45) or the cytosol of cultured *Arabidopsis* cells (46) circumvent several limitations. However, they often rely on non-endogenous baits (TRXs or GRXs from other cell compartments or species, mutated YAP1 from yeast), which introduce bias since the selectivity between TRX-type proteins and their targets can differ profoundly based on electrostatic surface properties (22). A study combining *in vitro* TRX-trapping and *in vivo* monobromobimane labeling identified a total of 114 TRX-targets from germinating Medicago seeds, including a small number of abundant mitochondrial proteins, such as HSP70, β-subunit of the mitochondrial ATP-synthase, pyruvate decarboxylase, succinate dehydrogenase, succinic semialdehyde dehydrogenase and peroxiredoxin IIF (13).

Here we establish isolated, actively respiring seedling mitochondria as an *in situ* model to detect and quantify specifically those Cys-thiol switches that are operated in the mitochondria in response to a redox transition that mimics the reactivation occurring during early seed germination **(Fig. 3A)**. The application of this model system to germination is further validated by the previous detection of the large majority of the mitochondrial proteins with operational thiol switches in seeds (**Fig. S10A**) (24). The reduction of target proteins was achieved by the endogenous thiol redox machinery, ensuring maximal target specificity. Thiol switching occurs specifically in the matrix of intact mitochondria, where NADPH is endogenously generated. The resulting enrichment for mitochondrial thiol-switched target proteins is illustrated by the comparison of the consensus subcellular localization of all identified proteins vs. those that showed thiol switching **(Fig. S7B)** (47). The reduction of target proteins is specific to the mitochondrial matrix where the full redox cascade can operate, as exemplified for respiratory Complex I, which showed thiol switching on matrix-exposed subunits, but not on the IMS exposed subunits **(Fig. 3F)**. Individual cases of reduction of non-matrix Cys are probably due to redox interaction between proteins that may occur during the short time window of the blocking step when mitochondria are ruptured. It is a strength of the differential iodoTMT-based approach that the individual cysteine thiol to undergo switching can be pinpointed and quantified in their degree of oxidation in most cases, whereas many other techniques only report relative changes. This provides a unique opportunity to compare the two approaches of fluorescent protein-based redox sensing and redox proteomics in the same system: in citrate-activated mitochondria roGFP2 underwent a reductive shift by Δ43.1 % as quantified by fluorimetry, compared to Δ30.4 % as quantified by MS/MS (only one of the two Cys-peptides involved in the roGFP disulfide was identified), indicating a qualitative and close to quantitative agreement between the two independent approaches.

Thiol switching can impact on the activity, stability, conformation and localisation of a protein, as well as its interaction with other proteins. Although activity of a switch, as demonstrated here, is necessary for those changes, it is not sufficient for a protein to be redox regulated (23). Several of the proteins we identify have been studied *in vitro*, through Cys mutagenesis and with a focus on the impact of Cys redox status on enzymatic activity. Fumarase and succinate dehydrogenase activity were inhibited by TRX-mediated reduction (28). Also for mitochondrial citrate synthase and isocitrate dehydrogenase redox regulation was observed (48, 49), while no change of activity was found for malate dehydrogenase 1 (50). Our study now provides the missing evidence for the operation of those thiol switches *in situ*, as a critical prerequisite for physiological redox regulation in mitochondria. The well-characterized thiol switch of the alternative oxidase (AOX) dimer was previously shown to respond to specific respiratory substrates supplied to isolated tobacco mitochondria, including citrate, but not others, including 2OG (51). Those observations elegantly match our findings at the proteome level and by *in situ* redox sensing. They further validate our interpretation that endogenous reduction of NADPH, but not NADH, by matrix metabolism leads to a re-start of the endogenous thiol redox systems in mitochondria. The thiol-switched Cys-peptide of the AOX1a, however, was not identified in our dataset, probably due to the cut-off criterion for the proteomic data evaluation (≥7 AA; the predicted Cys-peptide is 5 AA) **(Fig. S10B)**. While there is no prior evidence for Cys-based redox control of other respiratory chain components in plants, our observation of the respiratory chain as a hotspot of operational thiol switching is mirrored by thiol switches that regulate the activities of mitochondrial Complex I, Complex V in mammalian systems (recently reviewed in (23)).

A growing number of thiol redox proteomes has been generated for various organisms under resting and stress conditions, but none captures the transition between a quiescent and an active state of metabolism (37, 52–58). The global % oxidation of the inactive plant mitochondria (27.1 %) is higher than that found in previous studies of various organisms (within the range of 10 to 20 %); while metabolic activation established a % oxidation (13.1 %) that matched observations in other metabolically active systems. A noteworthy exception is a redox proteome of the heads and thoraces from fasting *Drosophila*, which showed 30.5 % oxidation overall (shifted from 19.6 % in the non-fasting control), i.e. a similar value as the quiescent plant mitochondria (57). Overexpression of catalase to scavenge H_2_O_2_ as a potential oxidant did not decrease the degree of oxidation in the fasting flies, indicating a dominant effect of the reductant supply through metabolism. Fasting *Drosophila* and quiescent plant mitochondria may have in common that metabolic flux is insufficient to provide enough NADPH to operate the thiol redox machinery effectively. By providing substrate to mitochondrial metabolism, the degree of Cys oxidation is effectively lowered, reaching comparable levels to the *in vivo* values of different metabolically active organisms.

We conclude that redox regulation contributes to efficient metabolism during early seed germination. To investigate if this strategy is conserved as a principle of metabolic regulation it will be interesting to look at other systems that show similarly pronounced metabolic transitions in the future; for instance, the metabolic activation of an egg cell after fertilization. That would make Cys-based redox regulation of chloroplast metabolism, which is the founding example and particularly well studied (59), a specific case of a more fundamental principle of regulation during metabolic transitions. Of further interest will be the role of mitochondrial thiol switching in seed dormancy. Since re-activation of energy and redox metabolism occurs directly at imbibition, it is plausible that this initial metabolic response influences the outcome of seed imbibition, ultimately executed by genetic programs under hormonal control. Under natural conditions a seed may undergo several imbibition–desiccation cycles during which dormancy levels are continuously readjusted by environmental factors. This raises the questions if there is a trade-off between dormancy and the protection of stored energy resources, and ultimately germination vigor (60), and to what extent thiol switching moderates this trade-off by providing an early level of control.

## MATERIALS AND METHODS

### Plant material and growth

Synchronized plants were used for seed production. Non-dormant and not-stratified seeds of *Arabidopsis thaliana* (L.) Heynh. Columbia-0 were used for all assays. Cytosolic ATeam 1.03 nD/nA (15) and mitochondrial roGFP2(-Grx1) (18, 19) lines were used for fluorimetry and confocal imaging; the latter also for redox proteomics. T-DNA insertion lines: *ntr a* x *ntr b* (27), SALK_539152 and SALK_545978, referred to as *ntr a/b* in this manuscript; *trx-o1* (28), SALK_042792; *gr2 epc-2* (20), SALK_040170 complemented for plastidic GR2 under control of the endogenous promotor, referred to as *gr2* in this manuscript. Seeds were placed on half-strength Murashige and Skoog medium supplemented with 10 mM 2-(N-morpholino)ethanesulfonic acid, pH 5.8 with KOH, 1 % (w/v) agar with or without 1 % (w/v) sucrose for monitoring the germination efficiency. Plants were cultivated under long-day conditions (16 h 75–100 μmol photons m^-2^s^-1^ at 22 °C, 8 h dark at 18 °C). For hydroponic plant cultures, conditions were as described before (15).

### Fluorimetry and confocal imaging

Fluorimetry and confocal imaging of fluorescent biosensors was performed as described (15, 61). For confocal imaging of isolated *Arabidopsis* embryos, seeds were imbibed on wet filter paper for 10-240 min before testa and endosperm were removed with tweezers. For fluorimetry of intact seeds during imbibition, seeds were placed in 384 well microplates. Rehydration was started during the measurement by injection of water by the built-in reagent injectors of a CLARIOstar plate reader (BMG LABTECH GmbH), while maintaining a constant temperature of 25 °C. Oxygen consumption of intact seeds was measured with MitoXpress Xtra fluorescent dye (Agilent) in a multi-well plate format as described (62).

### Metabolite measurements

Equal quantities of seeds from two individual plants were combined for each biological replicate. Extraction and quantification of metabolites by UPLC or GC-MS was performed as described (63–65) and normalized to seed dry weight. Dry seeds were imbibed at 25 °C for 0 h, 1 h or 4 h in unsealed reaction tubes with 250 μL H_2_O at 25 °C under gentle shaking before metabolite extraction. For extraction of organic acids, the water used for imbibition was analyzed separately to capture leached metabolites and the values for eluted organic acids in the aqueous fraction were added to the values obtained for seed extracts.

### Mitochondrial respiration assays

Functional mitochondria were isolated from *Arabidopsis* seedlings as recently described (15). For fluorimetry, respiration assays and redox proteomics, mitochondria were incubated in basic incubation medium (0.3 M sucrose, 10 mM NaCl, 2 mM MgSO_4_, 5 mM KH_2_PO_4_ and 10 mM TES, pH 7.5 with KOH). Oxygen consumption in respiratory state II was measured as described (66). For redox proteomics, mitochondrial fractions (~200 μg) were supplemented with 10 mM citrate or 10 mM 2OG for 25 min at 25 °C in a running unsealed Clark-type oxygen electrode keeping the medium aerated by stirring at 100 rpm as validated by online oxygen monitoring. Samples for quiescence were prepared from the identical preparation, also incubated for 25 min but without substrate addition.

### Differential iodoTMT labeling

Respiring and quiescent mitochondrial fractions were pelleted by centrifugation (30 s, 13000 *g*) and solubilized in iodoTMT-labeling buffer (8 M urea, 20 mM HEPES, pH 8.0, 1 mM EDTA, 0.01 % (v/v) Triton-X-100, 1 vial iodoTMT per replicate (ThermoFisher Scientific) to a final protein amount of ~200 μg and were incubated for 1.5 h at 30 °C in the dark, gently shaking. Insoluble proteins were removed by centrifugation (10 min, 18000 *g*), followed by acetone precipitation overnight (4-fold volume of pre-chilled 100 % (v/v) acetone, precipitation at −20 °C). Centrifugation of samples performed at room temperature (10 min, 18000 *g*). Protein pellets were washed twice with 80 % (v/v) and once with 100 % (v/v) acetone (proteins pelleted by centrifugation for 5 min at 18000 *g*). For differential iodoTMT labeling, proteins were resuspended in second labeling buffer (8 M urea, 20 mM HEPES, pH 8.0, 1 mM TCEP, 1 vial iodoTMT per replicate) to a final amount of 175 μg and were incubated for 1.5 h at 30 °C in the dark, while shaking gently. Samples were multiplexed with respect to their mass-tags (labels were swapped between individual replicates to compensate for hypothetical label biases). Unbound iodoTMTs were removed by overnight acetone precipitation (as for 1^st^ labeling) and dry protein pellets were used for trypsination.

### Trypsination, Immunoprecipitation and MS sample preparation

Proteins were dissolved in ABC buffer (25 mM ammonium bicarbonate, pH 7–8; 1 μL of buffer per 1 μg of protein). Trypsin was added in a protein to trypsin ratio of 100:1. Proteins were trypsinated at 37 °C for 2 h, gently shaking. Again, trypsin was added to a final protein to trypsin ratio of 50:1. After overnight incubation at 37 °C, 1/20 volume of 20 % (v/v) TFA was added to stop trypsinization and samples were centrifuged (10 min, 18000 *g*). Samples were cleared of TFA and ABC buffer by freeze-drying. Peptides were dissolved in TBS buffer (50 mM Tris-HCl, pH 7.6, 150 mM NaCl; 1 μL of buffer per 1 μg of peptides) and sonicated for 15 s. Insoluble particles were pelleted by centrifugation (10 min, 18000 *g*). For 350 μg of peptides, 350 μL of antibody resin was used (ThermoFisher Scientific). Before usage, antibody resin was washed three times with TBS and the slurry volume was replaced by addition of 175 μL TBS. IodoTMT-labeled Cys-peptides were bound to the antibody resin during overnight incubation with overhead rotation at 4 °C. Unbound peptides were removed by centrifugation (2 min, 1500 *g*). The antibody resin was incubated for 30 min with 1 mL TBS, gently shaking. After centrifugation (1 min, 1500 *g*), the antibody resin was washed three times with 1 mL TBS and two times with H_2_O. IodoTMT elution solution was added (400 μL, ThermoFisher Scientific) and incubated for 10 min with gentle agitation. After centrifugation (2 min, 1500 *g*), the supernatants were transferred to new reaction tubes and the elution solution was removed by freeze-drying. Peptides were dissolved in 20 μL of 5 % (v/v) acetonitrile and 0.1 % (v/v) TFA. After short centrifugation, 20 μL of 0.1 % (v/v) TFA was added. Samples were loaded on non-polar Pierce C18 StageTips (ThermoFisher Scientific), which were activated prior to the loading by addition of 20 μL of 100 % (v/v) acetonitrile. The C18-material was equilibrated with 20 μL of 0.1 % (v/v) TFA. Bound peptides were washed with 20 μL of 0.1 % (v/v) TFA and were eluted by addition of 40 μL of 60 % (v/v) acetonitrile and 0.1 % (v/v) TFA. Eluted peptides were placed in a centrifugal evaporator to remove the acetonitrile/TFA solution. For mass spectrometry analysis, peptides were finally dissolved in 20 μL of 5 % (v/v) acetonitrile and 0.1 % (v/v) acetic acid.

### LC-MS/MS-analysis and raw data processing

Peptide samples were analyzed by LC-MS/MS as described recently (67), with specific modifications. In brief, iodoTMT samples were loaded onto the LC system with 8 μL of 0.1% (v/v) acetic acid at a constant flow rate of 500 nL/min. Peptides were eluted using a non-linear 180 min gradient from 1% to 99% of 0.1% (v/v) acetic acid in acetonitrile at a flow rate of 300 nL/min and directly injected into an Orbitrap Velos Pro (Thermo Scientific). Each of the samples was analyzed in two technical replicates. Database searching and quantification was performed within the framework of the MaxQuant software suite (1.5.5.1) (68) and spectra were searched against the *Arabidopsis thaliana* TAIR10 database, which was complemented with the sequence for the roGFP2-Grx1 sensor protein. The percentage of Cys-oxidation (% ox) for the razor peptides was calculated from the reporter ion intensity of the 2^nd^ label and the sum of the reporter ion intensities of the 1^st^ and the 2^nd^ label.

## ACKNOWLEDGEMENTS

We thank Jürgen Eirich (Münster) for support with proteomic data handling and the Deutsche Forschungsgemeinschaft (DFG) for financial support through the Emmy-Noether program (SCHW1719/ 1-1), the priority program SPP1710 ‘Dynamics of thiol-based redox switches in cellular physiology’ (SCHW1719/ 7-1, ME1567/9-1/2, LI 984/3-1/2), the infrastructure grant INST 211/744-1 FUGG and the project grants (SCHW1719/5-1, FI1655/3-1) as part of the package PAK918.

## REFERENCES

1. Sallon S, et al. (2008) Germination, genetics, and growth of an ancient date seed. Science 320(5882):1464–1464.

2. Pracharoenwattana I, Cornah JE, & Smith SM (2005) Arabidopsis peroxisomal citrate synthase is required for fatty acid respiration and seed germination. Plant Cell 17(7):2037–2048.

3. Eastmond PJ & Graham IA (2001) Re-examining the role of the glyoxylate cycle in oilseeds. Trends Plant Sci 6(2):72–77.

4. Galland M, et al. (2014) Dynamic proteomics emphasizes the importance of selective mRNA translation and protein turnover during Arabidopsis seed germination. Mol Cell Proteomics 13(1):252–268.

5. Nee G, Xiang Y, & Soppe WJJ (2017) The release of dormancy, a wake-up call for seeds to germinate. Current opinion in plant biology 35:8–14.

6. El-Maarouf-Bouteau H & Bailly C (2008) Oxidative signaling in seed germination and dormancy. Plant signaling & behavior 3(3):175–182.

7. Arc E, et al. (2011) Reboot the system thanks to protein post-translational modifications and proteome diversity: How quiescent seeds restart their metabolism to prepare seedling establishment. Proteomics 11(9):1606–1618.

8. Buchanan BB (2017) The path to thioredoxin and redox regulation beyond chloroplasts. Plant Cell Physiol 58(11):1826–1832.

9. Meyer Y, Belin C, Delorme-Hinoux V, Reichheld JP, & Riondet C (2012) Thioredoxin and glutaredoxin systems in plants: molecular mechanisms, crosstalks, and functional significance. Antioxidants & redox signaling 17(8):1124–1160.

10. Hopkins FG & Morgan EJ (1943) Appearance of glutathione during the early stages of the germination of seeds. Nature 152:288–290.

11. Kobrehel K, et al. (1992) Specific reduction of wheat storage proteins by thioredoxin-h. Plant Physiol 99(3):919–924.

12. Yano H, Wong JH, Cho MJ, & Buchanan BB (2001) Redox changes accompanying the degradation of seed storage proteins in germinating rice. Plant Cell Physiol 42(8):879–883.

13. Alkhalfioui F, et al. (2007) Thioredoxin-linked proteins are reduced during germination of *Medicago truncatula* seeds. Plant Physiol 144(3):1559–1579.

14. Wong JH, et al. (2004) Thioredoxin reduction alters the solubility of proteins of wheat starchy endosperm: An early event in cereal germination. Plant Cell Physiol 45(4):407–415.

15. De Col V, et al. (2017) ATP sensing in living plant cells reveals tissue gradients and stress dynamics of energy physiology. eLife 6:e26770.

16. Nakabayashi K, Okamoto M, Koshiba T, Kamiya Y, & Nambara E (2005) Genome-wide profiling of stored mRNA in *Arabidopsis thaliana* seed germination: epigenetic and genetic regulation of transcription in seed. Plant J 41(5):697–709.

17. Shu K, Liu XD, Xie Q, & He ZH (2016) Two faces of one seed: Hormonal regulation of dormancy and germination. Mol Plant 9(1):34–45.

18. Albrecht SC, et al. (2014) Redesign of genetically encoded biosensors for monitoring mitochondrial redox status in a broad range of model eukaryotes. J Biomol Screen 19(3):379–386.

19. Schwarzländer M, et al. (2008) Confocal imaging of glutathione redox potential in living plant cells. J Microsc-Oxford 231(2):299–316.

20. Marty L, et al. (2019) Arabidopsis glutathione reductase 2 is indispensable in plastids, while mitochondrial glutathione is safeguarded by additional reduction and transport systems. bioRxiv:610477.

21. Fricker MD (2016) Quantitative redox imaging software. Antioxidants & redox signaling 24(13):752–762.

22. Berndt C, Schwenn JD, & Lillig CH (2015) The specificity of thioredoxins and glutaredoxins is determined by electrostatic and geometric complementarity. Chem Sci 6(12):7049–7058.

23. Nietzel T, Mostertz J, Hochgräfe F, & Schwarzländer M (2017) Redox regulation of mitochondrial proteins and proteomes by cysteine thiol switches. Mitochondrion 33.

24. Xiang Y, et al. (2016) Sequence Polymorphisms at the REDUCED DORMANCY5 Pseudophosphatase Underlie Natural Variation in Arabidopsis Dormancy. Plant Physiol 171(4):2659–2670.

25. Moller IM & Rasmusson AG (1998) The role of NADP in the mitochondrial matrix. Trends Plant Sci 3(1):21–27.

26. Reichheld JP, Meyer E, Khafif M, Bonnard G, & Meyer Y (2005) AtNTRB is the major mitochondrial thioredoxin reductase in *Arabidopsis thaliana*. Febs Lett 579(2):337–342.

27. Reichheld JP, et al. (2007) Inactivation of thioredoxin reductases reveals a complex interplay between thioredoxin and glutathione pathways in Arabidopsis development. Plant Cell 19(6):1851–1865.

28. Daloso DM, et al. (2015) Thioredoxin, a master regulator of the tricarboxylic acid cycle in plant mitochondria. P Natl Acad Sci USA 112(11):E1392–E1400.

29. Weerapana E, et al. (2010) Quantitative reactivity profiling predicts functional cysteines in proteomes. Nature 468(7325):790–795.

30. Bak DW, Pizzagalli MD, & Weerapana E (2017) Identifying functional cysteine residues in the mitochondria. ACS chemical biology.

31. Braun HP, et al. (2014) The life of plant mitochondrial complex I. Mitochondrion 19 Pt B:295–313.

32. Balmer Y, et al. (2004) Thioredoxin links redox to the regulation of fundamental processes of plant mitochondria. P Natl Acad Sci USA 101(8):2642–2647.

33. Rouhier N, et al. (2005) Identification of plant glutaredoxin targets. Antioxidants & redox signaling 7(7-8):919–929.

34. Yoshida K, Noguchi K, Motohashi K, & Hisabori T (2013) Systematic exploration of thioredoxin target proteins in plant mitochondria. Plant Cell Physiol 54(6):875–892.

35. Winger AM, Taylor NL, Heazlewood JL, Day DA, & Millar AH (2007) Identification of intra-and intermolecular disulfide bonding in the plant mitochondrial proteome by diagonal gel electrophoresis. Proteomics 7(22):4158–4170.

36. Marti MC, et al. (2009) Mitochondrial and nuclear localisation of a novel Pea thioredoxin: Identification of its mitochondrial target proteins. Plant Physiol 150(2):646–657.

37. Chouchani ET, et al. (2016) Mitochondrial ROS regulate thermogenic energy expenditure and sulfenylation of UCP1. Nature 532(7597):112–116.

38. Marty L, et al. (2009) The NADPH-dependent thioredoxin system constitutes a functional backup for cytosolic glutathione reductase in Arabidopsis. P Natl Acad Sci USA 106(22):9109–9114.

39. Hildebrandt TM, Nesi AN, Araujo WL, & Braun HP (2015) Amino acid catabolism in plants. Mol Plant 8(11):1563–1579.

40. Law SR, et al. (2012) Nucleotide and RNA metabolism prime translational initiation in the earliest events of mitochondrial biogenesis during Arabidopsis germination. Plant Physiol 158(4):1610–1627.

41. Raveneau MP, Benamar A, & Macherel D (2017) Water content, adenylate kinase, and mitochondria drive adenylate balance in dehydrating and imbibing seeds. J Exp Bot 68(13):3501–3512.

42. Paszkiewicz G, Gualberto JM, Benamar A, Macherel D, & Logan DC (2017) Arabidopsis seed mitochondria are bioenergetically active immediately upon imbibition and specialize via biogenesis in preparation for autotrophic growth. Plant Cell 29(1):109–128.

43. Hourmant A & Pradet A (1981) Oxidative-phosphorylation in germinating lettuce seeds *(Lactuca-Sativa)* during the 1st hours of imbibition. Plant Physiol 68(3):631–635.

44. Benamar A, Tallon C, & Macherel D (2003) Membrane integrity and oxidative properties of mitochondria isolated from imbibing pea seeds after priming or accelerated ageing. Seed Sci Res 13(1):35–45.

45. Engelhard J, et al. (2011) *In situ* kinetic trapping reveals a fingerprint of reversible protein thiol oxidation in the mitochondrial matrix. Free Radical Bio Med 50(10):1234–1241.

46. Waszczak C, et al. (2014) *Sulfenome* mining in *Arabidopsis thaliana*. P Natl Acad Sci USA 111(31):11545–11550.

47. Hooper CM, et al. (2014) SUBAcon: a consensus algorithm for unifying the subcellular localisation data of the Arabidopsis proteome. Bioinformatics 30(23):3356–3364.

48. Schmidtmann E, et al. (2014) Redox regulation of Arabidopsis mitochondrial citrate synthase. Mol Plant 7(1):156–169.

49. Yoshida K & Hisabori T (2014) Mitochondrial isocitrate dehydrogenase is inactivated upon oxidation and reactivated by thioredoxin-dependent reduction in Arabidopsis. Frontiers in Environmental Science.

50. Yoshida K & Hisabori T (2016) Adenine nucleotide-dependent and redox-independent control of mitochondrial malate dehydrogenase activity in *Arabidopsis thaliana*. Bba-Bioenergetics 1857(6):810–818.

51. Vanlerberghe GC, Day DA, Wiskich JT, Vanlerberghe AE, & Mcintosh L (1995) Alternative oxidase activity in tobacco leaf mitochondria - dependence on tricarboxylic-acid cycle-mediated redox regulation and pyruvate activation. Plant Physiol 109(2):353–361.

52. Leichert LI, et al. (2008) Quantifying changes in the thiol redox proteome upon oxidative stress *in vivo*. P Natl Acad Sci USA 105(24):8197–8202.

53. Brandes N, Reichmann D, Tienson H, Leichert LI, & Jakob U (2011) Using quantitative redox proteomics to dissect the yeast *redoxome*. J Biol Chem 286(48):41893–41903.

54. Brandes N, et al. (2013) Time line of redox events in ageing postmitotic cells. eLife 2:e00306.

55. Araki K, et al. (2016) Redox sensitivities of global cellular cysteine residues under reductive and oxidative stress. Journal of proteome research 15(8):2548–2559.

56. Knoefler D, et al. (2012) Quantitative *in vivo* redox sensors uncover oxidative stress as an early event in life. Mol Cell 47(5):767–776.

57. Menger KE, et al. (2015) Fasting, but not aging, dramatically alters the redox status of cysteine residues on proteins in *Drosophila melanogaster*. Cell Rep 11(12):1856–1865.

58. Topf U, et al. (2018) Quantitative proteomics identifies redox switches for global translation modulation by mitochondrially produced reactive oxygen species. Nat Commun 9.

59. Buchanan BB (2016) The path to thioredoxin and redox regulation in chloroplasts. Annual review of plant biology 67:1–24.

60. Nguyen TP, Keizer P, van Eeuwijk F, Smeekens S, & Bentsink L (2012) Natural variation for seed longevity and seed dormancy are negatively correlated in Arabidopsis. Plant Physiol 160(4):2083–2092.

61. Nietzel T, et al. (2019) The fluorescent protein sensor roGFP2-Orp1 monitors *in vivo* H_2_O_2_ and thiol redox integration and elucidates intracellular H_2_O_2_ dynamics during elicitor-induced oxidative burst in Arabidopsis. New Phytol 221(3):1649–1664.

62. Sechet J, et al. (2015) The ABA-deficiency suppressor locus HAS2 encodes the PPR protein LOI1/MEF11 involved in mitochondrial RNA editing. Mol Plant 8(4):644–656.

63. Bürstenbinder K, et al. (2010) Inhibition of 5 ‘-methylthioadenosine metabolism in the Yang cycle alters polyamine levels, and impairs seedling growth and reproduction in Arabidopsis. Plant J 62(6):977–988.

64. Abu el Maaty MA, et al. (2018) Activation of pro-survival metabolic networks by 1,25(OH)_2_D_3_ does not hamper the sensitivity of breast cancer cells to chemotherapeutics. Cancer Metab 6.

65. Weger M, et al. (2018) Glucocorticoid deficiency causes transcriptional and post-transcriptional reprogramming of glutamine metabolism. Ebiomedicine 36:376–389.

66. Sweetlove LJ, Taylor NL, & Leaver CJ (2007) Isolation of intact, functional mitochondria from the model plant *Arabidopsis thaliana*. Methods in Molecular Biology 372:125–136.

67. Sievers S, et al. (2018) Comprehensive redox profiling of the thiol proteome of *Clostridium difficile*. Mol Cell Proteomics 17(5):1035–1046.

68. Cox J & Mann M (2008) MaxQuant enables high peptide identification rates, individualized p.p.b.-range mass accuracies and proteome-wide protein quantification. Nat Biotechnol 26(12):1367–1372.

