## Supplementary material for "Redox-mediated Kick-Start of Mitochondrial Energy Metabolism drives Resource-efficient Seed Germination": SI Figures and Table

### Supplemental Figures

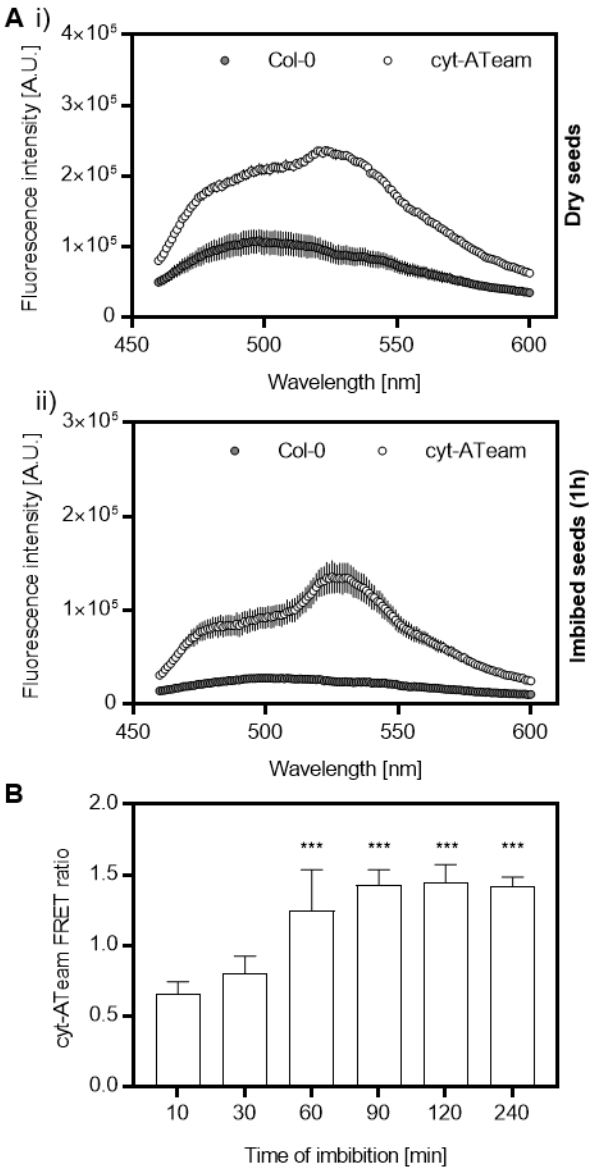


**Fig. S1. Cytosolic $MgATP^{2-}$ dynamics monitored by ATeam in *Arabidopsis* seeds.** (A) Emission scans performed for Col-0 seeds and for cytosolic ATeam seeds (cyt-ATeam) before (i) and after 1 h (ii) of addition of water to start imbibition (mean ± SD; *n* = 3-4 seed batches). (B) Cytosolic $MgATP^{2-}$ in seeds was measured with the fluorescent biosensor cyt-ATeam in response to imbibition by confocal imaging. Testa and endosperm were removed from intact embryos after imbibition for 10 to 240 min. The radicles of isolated embryos were analyzed and FRET ratios are plotted (mean + SD; *n* = 8-13 embryos; ***$p < 0.001$ with two-sided Student's *t*-test, if compared to 10 min of imbibition).

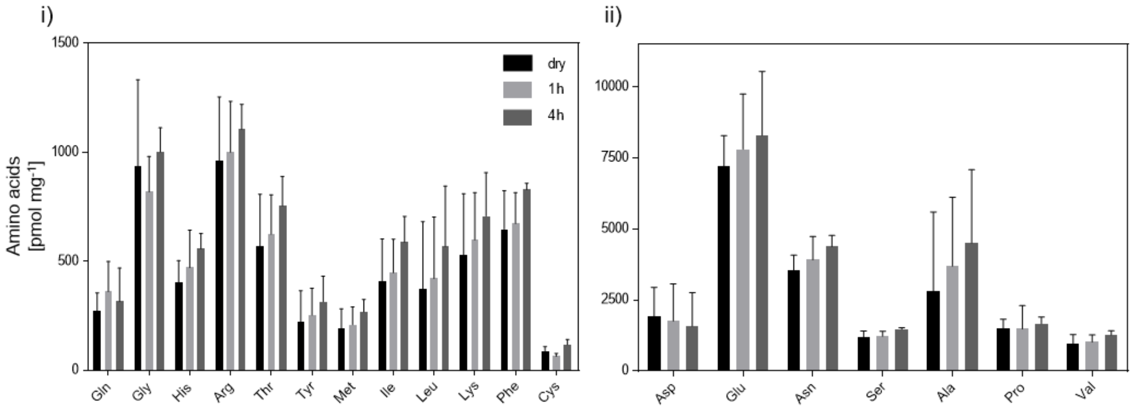


**Fig. S2. Amino acid profiles of dry and imbibed *Arabidopsis* seeds.** Free amino acids in total seed extracts of dry seeds or after 1 and 4 h of imbibition (*n* = 4; mean normalized to seed dry weight + SD, no statistical significant difference according to two-way ANOVA and Tukey`s multiple comparison test). Free cysteine was quantified after derivatization with monobromobimane in a separate extraction and analytics. **i**) Lowly abundant amino acids and **ii**) highly abundant amino acids.

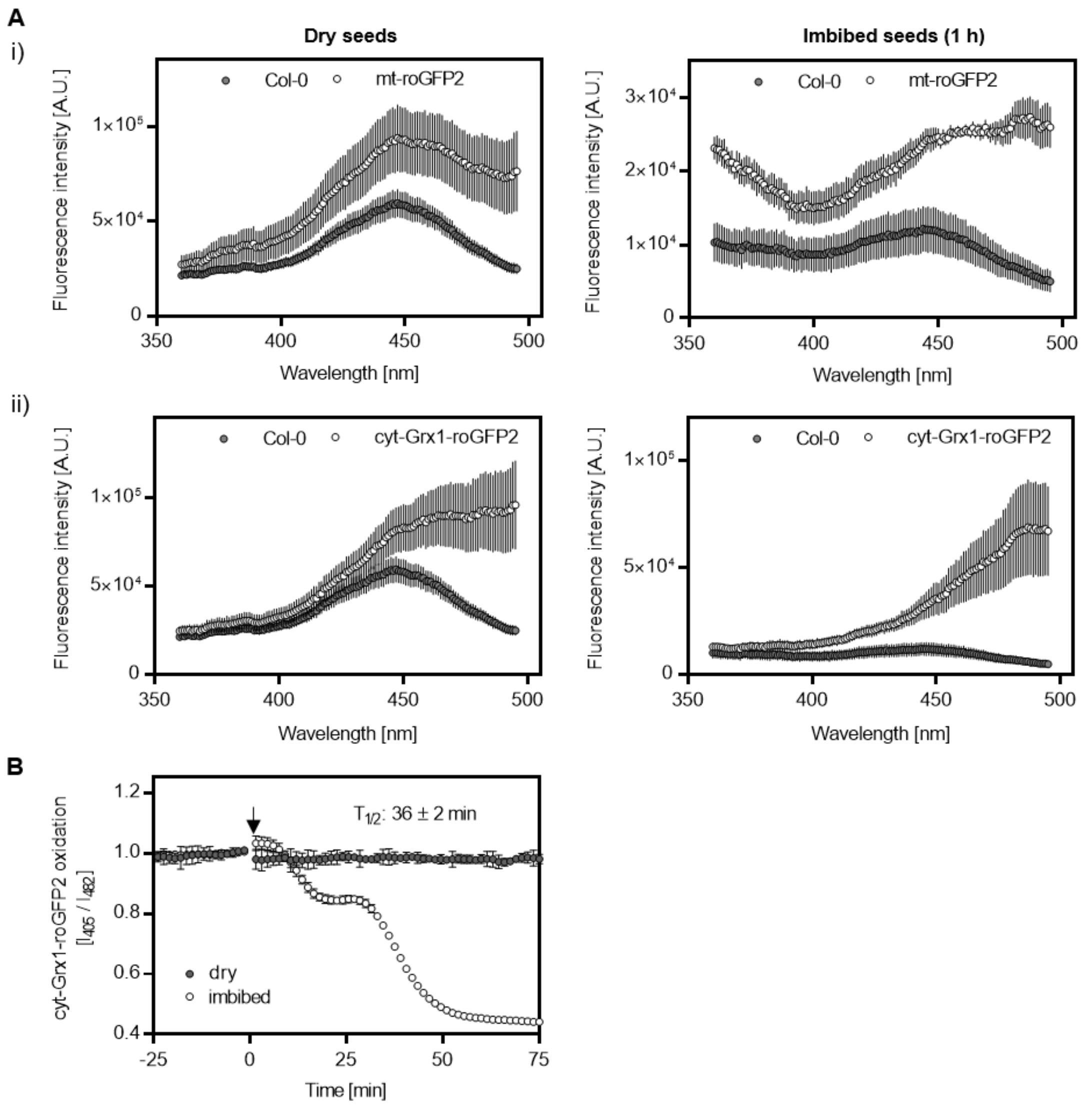


**Fig. S3. $E_{GSH}$ dynamics in the mitochondrial matrix and the cytosol measured by roGFP2 fluorescence in *Arabidopsis* seeds.** (**A**) Excitation spectra of *Arabidopsis* seeds expressing roGFP2-based probes in the mitochondrial matrix (**i**, mt-roGFP2) and the cytosol (**ii**, cyt-Grx1-roGFP2). Spectra recorded for dry seeds (left panels) and after 1 h imbibition (right panels) of Col-0 seeds and both sensor lines (mean ± SD; *n* = 3-4 seed batches). (**B**) Cytosolic Grx1-roGFP2 dynamics at imbibition of intact seeds monitored with a plate reader-based setup (as in **Fig. 1 A** and **B**). Arrow indicates addition of water, $T_{1/2}$ gives time point of half-maximal sensor response (*n* = 6 seed batches; mean normalized to last value before injection of water ± SD and corrected for Col-0 auto-fluorescence).

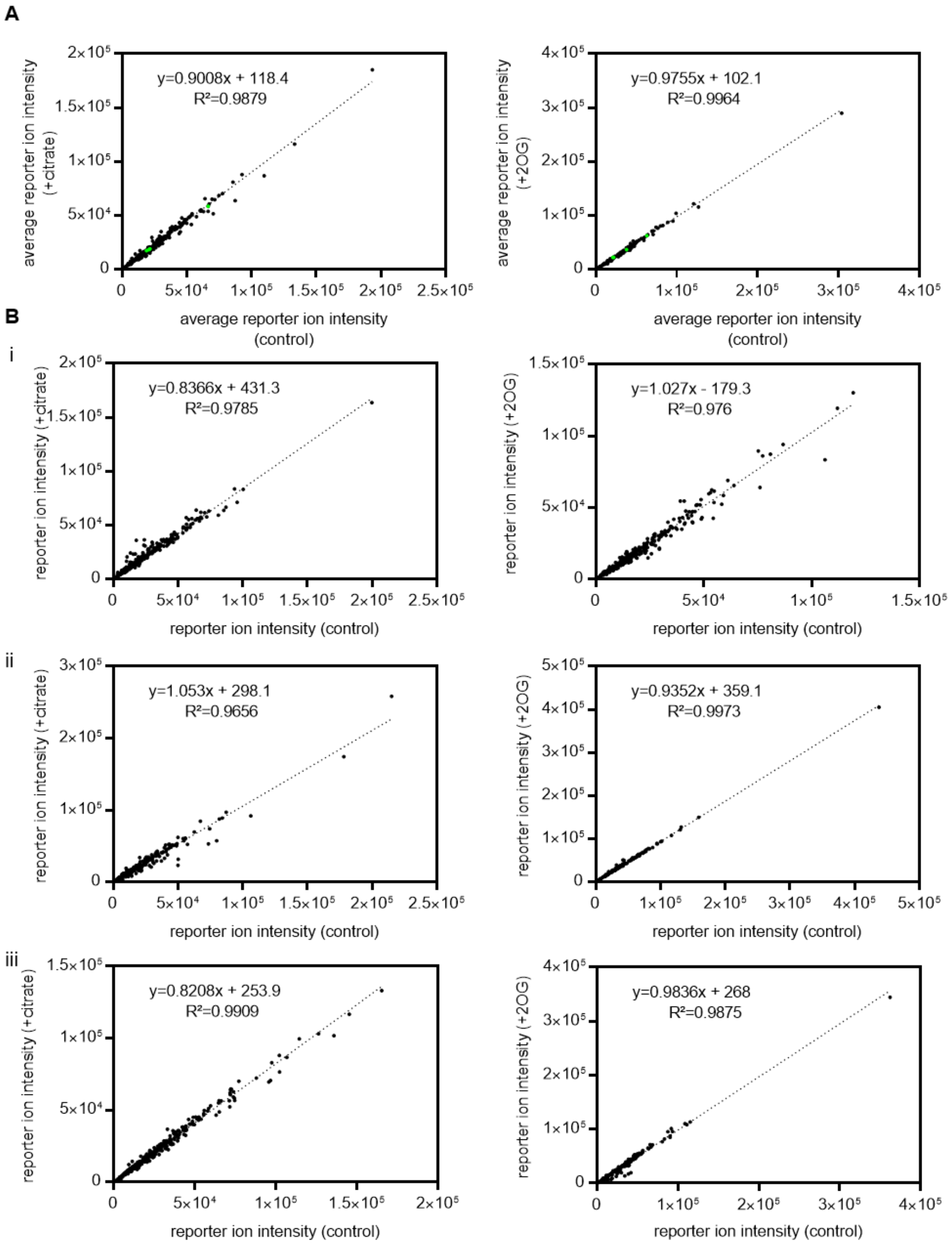


**Fig. S4. Reporter ion intensities between Cys-peptides of the control and the treated mitochondrial fractions.** (**A**) Average reporter ion intensity of all identified Cys-peptides ($n$ = 3) plotted for non-treated mitochondria after 25 min and for those after substrate application (left: 10 mM citrate; right: 10 mM 2OG). Reporter ion intensity of 1$^{st}$ and 2$^{nd}$ tag added together as a proxy for peptide abundance. Linear regression (dashed lines with corresponding regression equations) were fitted to the reporter ion intensities. In green, Cys-peptides of the mitochondrial localized roGFP2-Grx1, accounting for approximately 2 % of the cumulated overall reporter ion intensities (for all Cys-peptides identified with $n$ = 3). (**B**) As in (**A**), only for individual replicates (**i**, **ii** & **iii**).

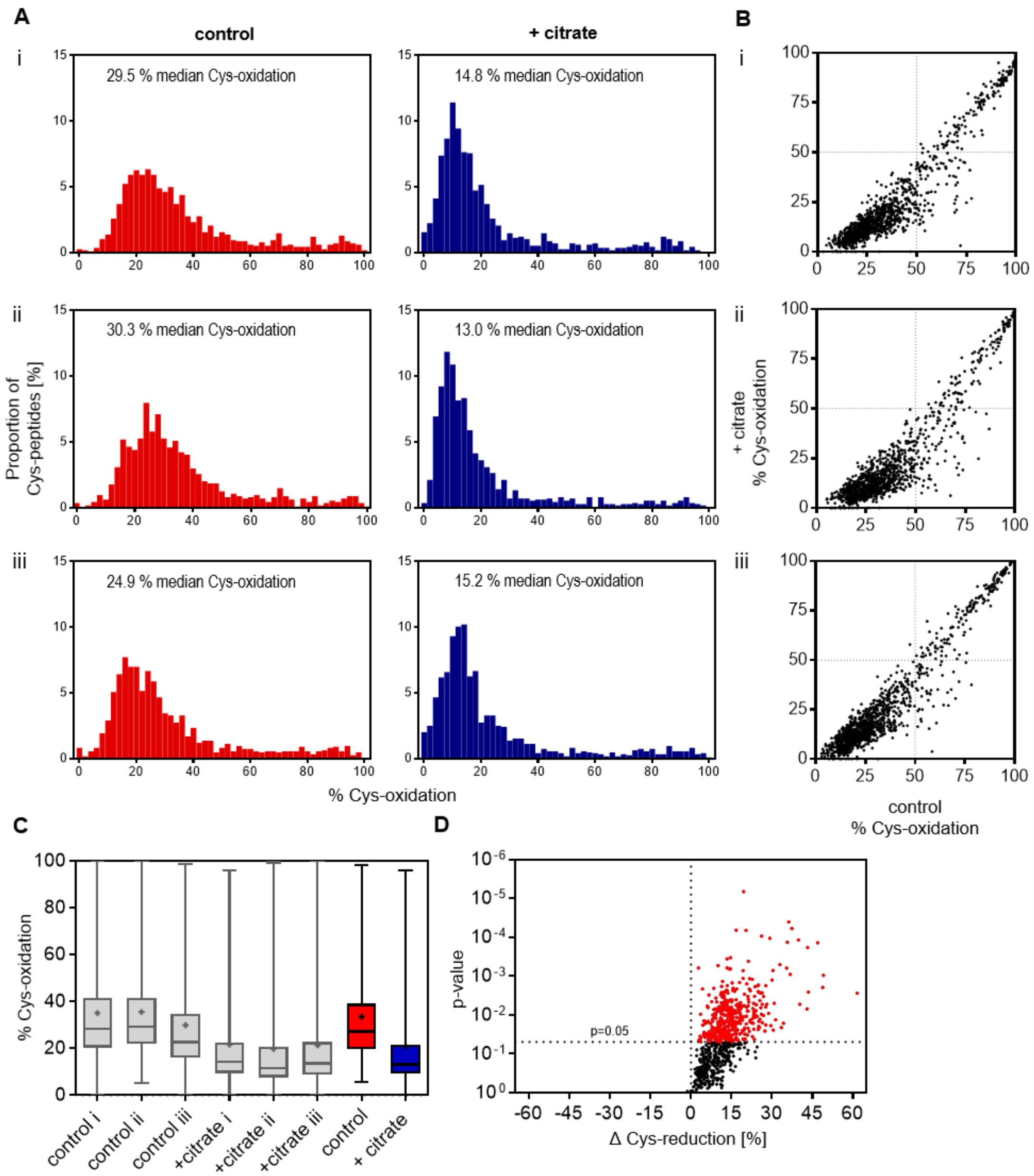


**Fig. S5. The redox landscape of Cys-peptides of quiescent and citrate-respiring mitochondria.** (**A**) Mitochondrial fractions of *Arabidopsis* seedlings incubated for 25 min with 10 mM citrate. Redox states of individual peptides quantified with an iodoTMT-based MS/MS approach for control mitochondrial fractions (red bars) or after citrate addition (blue bars). The distribution of cysteine-peptide oxidation levels is given for the replicates i, ii & iii; the proportion of the total number of peptides in each 2 % quantile of percentage oxidation is plotted. (**B**) The degree of Cys-oxidation of control mitochondrial fractions is plotted against the degree of Cys-oxidation after citrate addition for individual Cys-peptides. (**C**) Box-plots of the Cys-oxidation state (in percent of total oxidation) are given for all replicates (grey box-plots) and for their averages (red & blue box-plots; means indicated by crosses). (**D**) The *p*-values (two-sided Student`s *t*-test) of individual Cys-peptides are plotted against their Δ Cys-reduction levels (at citrate application) in a volcano-plot; in red all *p*-values $< 0.05$.

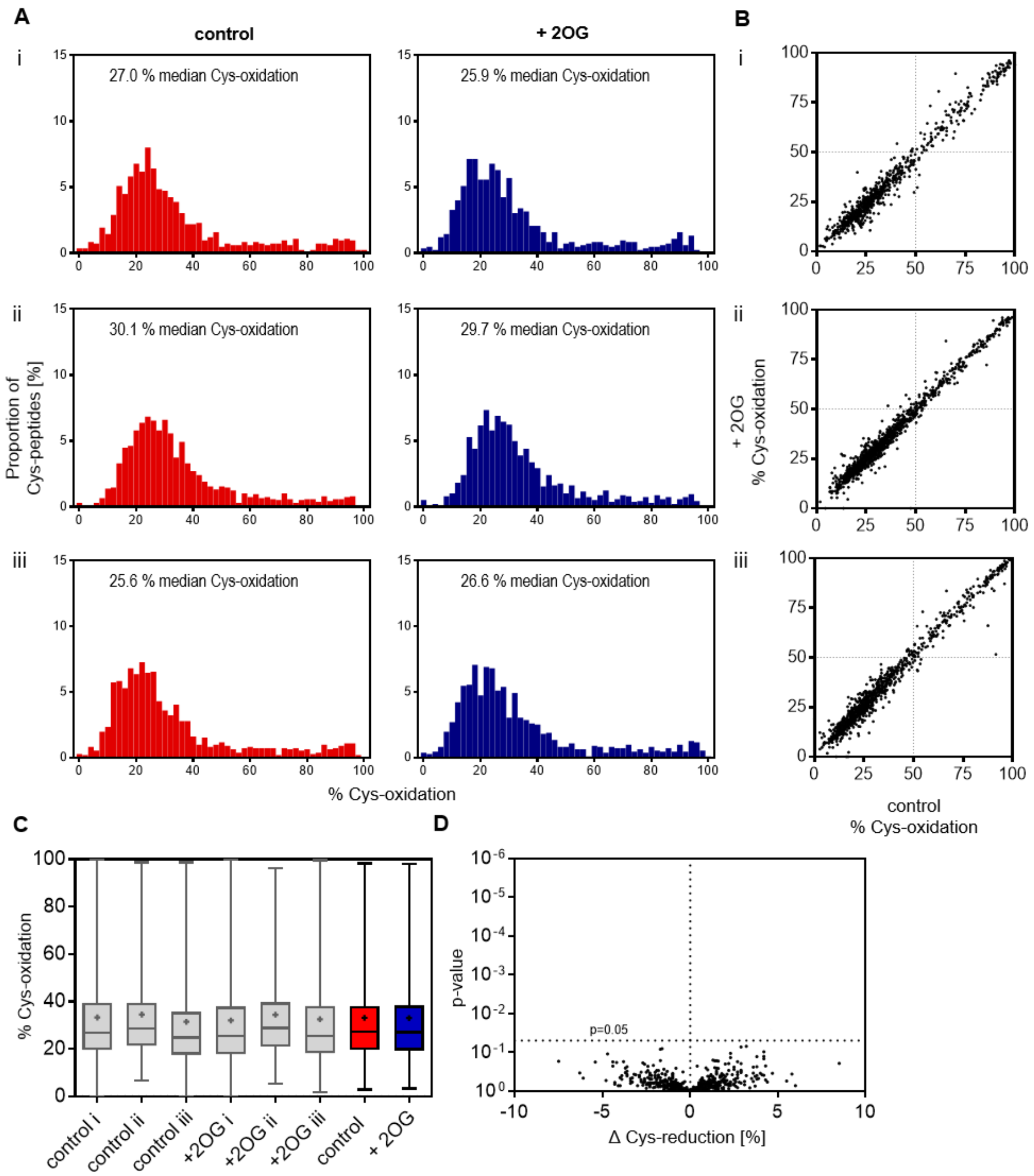


**Fig. S6. The redox landscape of Cys-peptides of quiescent and 2OG-respiring mitochondria.** (**A**) Mitochondrial fractions of *Arabidopsis* seedlings incubated for 25 min with 10 mM 2-oxoglutarate (2OG). Redox states of individual peptides quantified with an iodoTMT-based MS/MS approach for control mitochondrial fractions (red bars) or after 2OG addition (blue bars). The distribution of cysteine-peptide oxidation levels is given for the replicates i, ii & iii; the proportion of the total number of peptides in each 2 % quantile of percentage oxidation is plotted. (**B**) The degree of Cys-oxidation of control mitochondrial fractions is plotted against the degree of Cys-oxidation after 2OG addition for individual Cys-peptides. (**C**) Box-plots of the Cys-oxidation state (in percent of total oxidation) are given for all replicates (grey box-plots) and for their averages (red & blue box-plots; means indicated by crosses). (**D**) The *p*-values (two-sided Student`s *t*-test) of individual Cys-peptides are plotted against their Δ Cys-reduction levels (at 2OG application) in a volcano-plot; in red all *p*-values $< 0.05$.

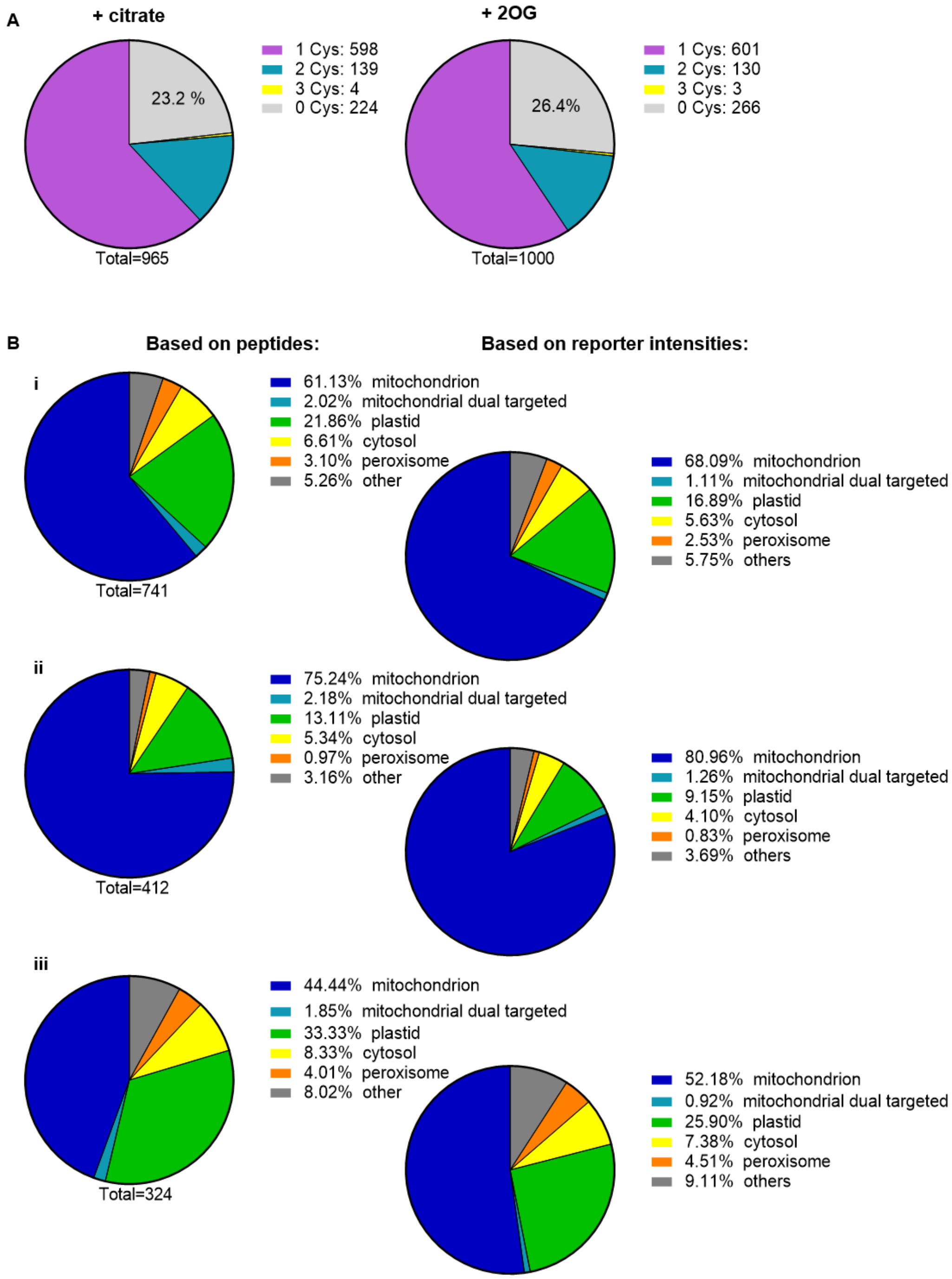


**Fig. S7. Quantitative characteristics of the Cys-peptides at the proteome level.** (**A**) Number of Cys-residues per peptide for all identified Cys-peptides (*n* = 3 for both respiratory conditions) in the citrate-supplemented mitochondrial fractions (left) and those supplemented with 2OG (right). (**B**) Annotated subcellular localization of all (**i**) identified Cys-peptides (*n* = 3) based on the *Arabidopsis* subcellular localization database (SUBA3, consensus localization) (47), either on the basis of number of identified peptides per cellular compartment (left) or on the corresponding reporter ion intensities (right). (**ii**) The annotated subcellular localization for the subset of significantly reduced Cys-peptides (*t*-test corrected for multiple comparisons by Benjamini, Krieger & Yekutieli with < 2 % FDR) and (**iii**) for non-redox-shifted Cys-peptides.

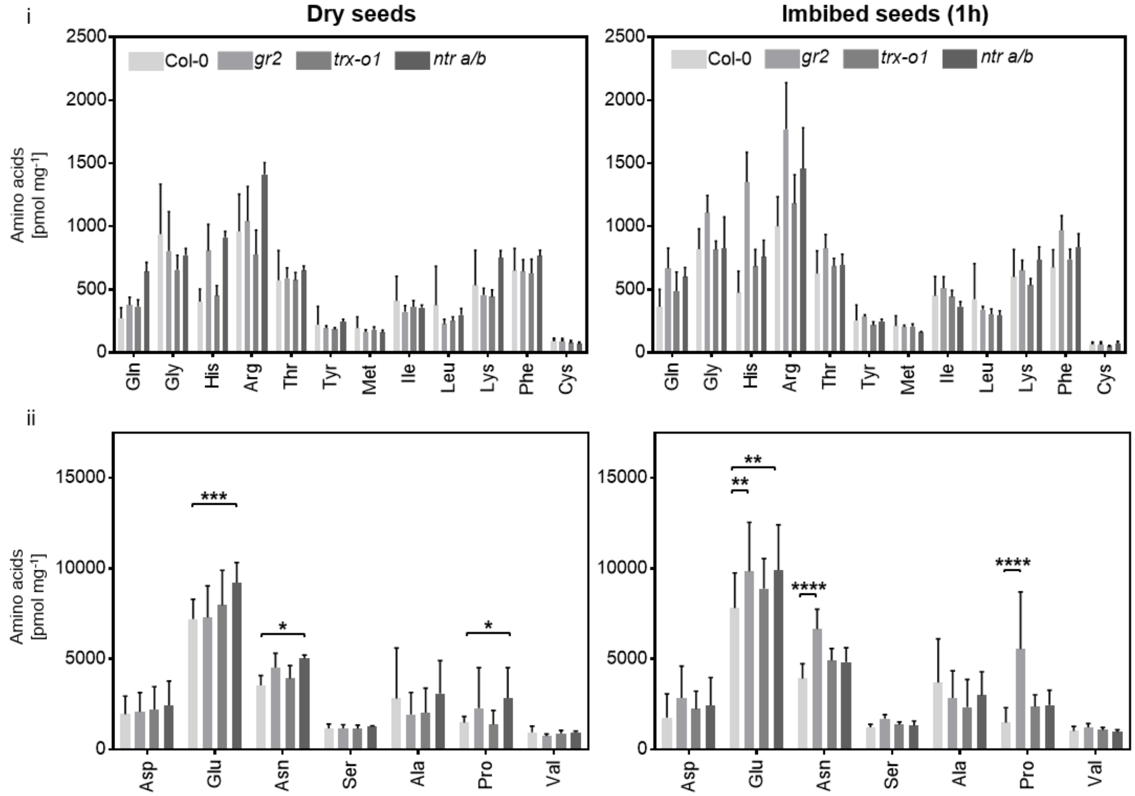


**Fig. S8. Amino acid profiles of dry and imbibed *Arabidopsis* seeds with impaired thiol redox machinery.** Free amino acid concentrations in total seed extracts before and after 1 h imbibition. Lowly abundant amino acids (**i**), highly abundant amino acids (**ii**) ($n$ = 4; mean normalized to seed dry weight + SD). Free cysteine was quantified after derivatization with monobromobimane by independent extraction and analysis. Significantly different values of the mutants in comparison to the Col-0 seeds are indicated by asterisks (two-way ANOVA and Dunnett`s multiple comparison test, *$p < 0.05$, **$p < 0.01$, ***$p < 0.001$, ****$p < 0.0001$).

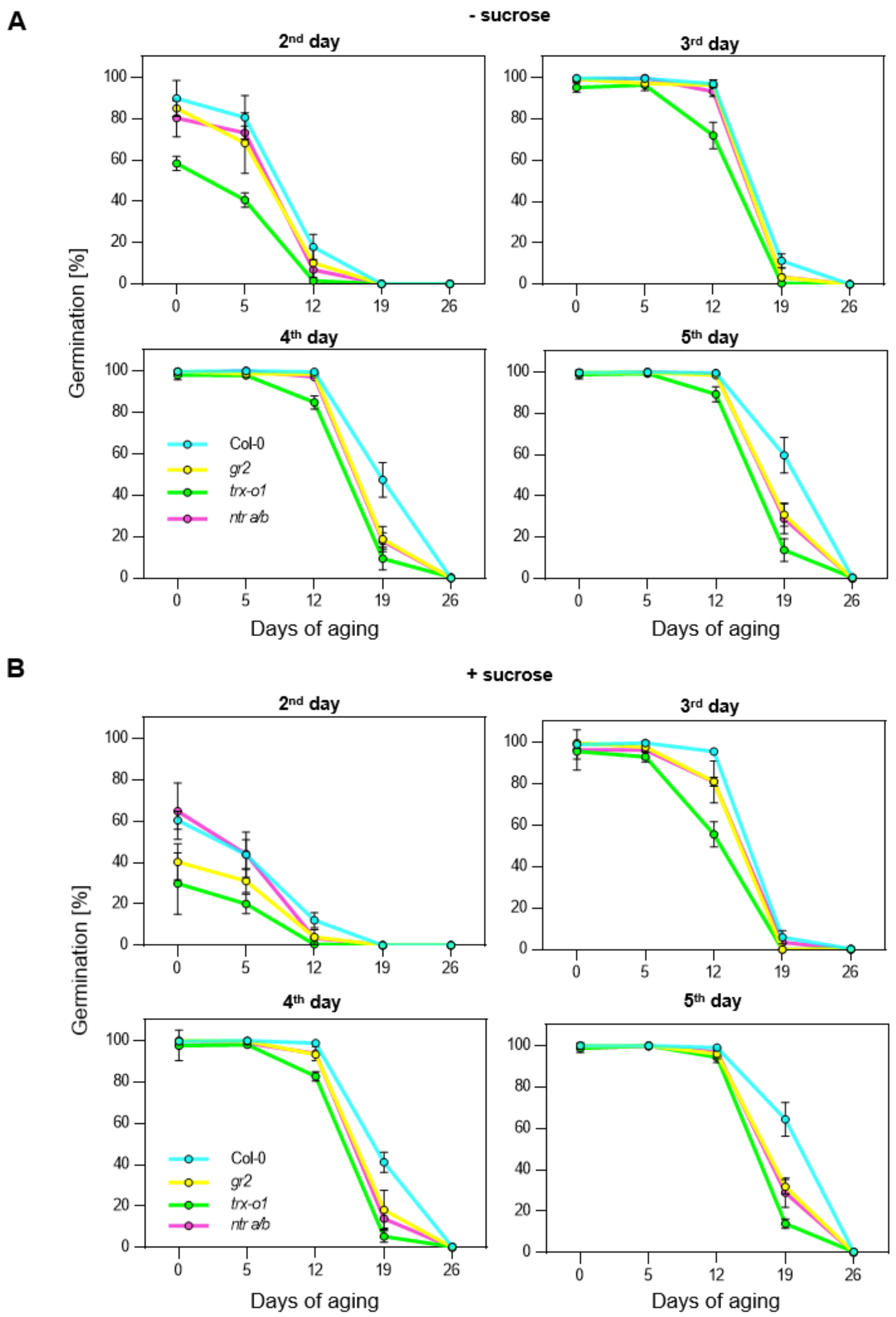


**Fig. S9. Germination of *Arabidopsis* seeds impaired in the thiol redox machinery after controlled deterioration.** *Arabidopsis* seeds were stored for 0, 5, 12, 19 and 26 days at 37 °C and 75 % rel. humidity. Seeds incubated on plates (0.5x MS media, 10 mM MES, pH 5.8 with KOH, 0.8 % (w/v) phytagel). (**A**) Media without sucrose and (**B**) with 1 % (w/v) sucrose. Radicle penetration of the testa and the endosperm was used as the criterion of germination. Germination was tracked for days 2-5 after sowing (mean ± SD, *n* = 200-250). Statistically different germination rates of *gr2*, *trx-o1* and *ntr a/b* as compared to Col-0 seeds are highlighted in **Table S1**.

**A**

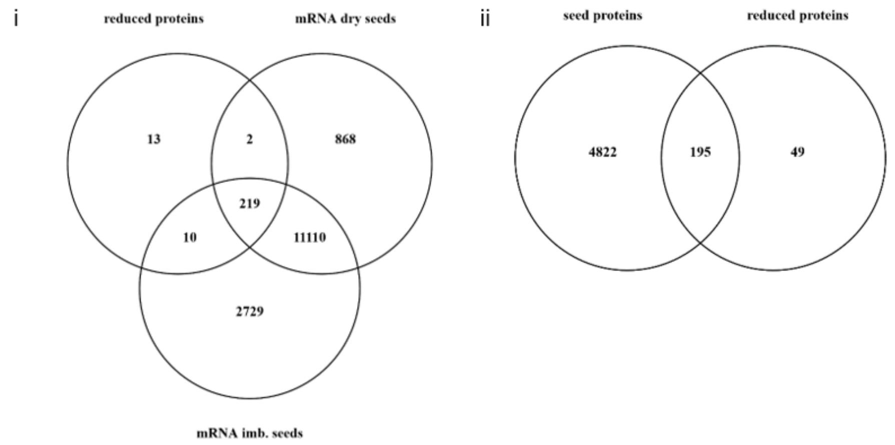


**B**

```
AOX2        AKMKITRKDGSDWPWNCFMPWETYQANLSIDLKKHHVPKNIADKVAYRIVKLLRIPTDIF
AOX1d       --TKITKPDGSAWKWNCFQPWDSYKPDVSIDVTKHHKPSNFTDKFAYWTVQTLKIPVQLF
AOX1a       --NKITKEDGSEWKWNCFRPWETYKADITIDLKKHHVPTTFLDRIAYWTVKSLRWPTDLF
AOX1b       --MKITKEDGTEWKWSCFRPWETYKSDLTIDLKKHHVPSTLPDKLAYWTVKSLRWPTDLF
AOX1c       --MKITKEDGTEWKWSCFRPWETYKADLTIDLKKHHVPSTLPDKIAYWMVKSLRWPTDLF
               ***: **: * *.** **::*: :::**:.*** *..: *:.**  *: *: *.::*
```

**Fig. S10. Validation of the model system and the Cys redox proteomic approach.** (**A**) Presence of proteinaceous cysteinyl redox switches identified in the purified mitochondria model in seeds. i) proteins with operational Cys-based redox switches for which also mRNA was identified in dry and/or imbibed seeds (16). Exclusively identified in the group of reduced proteins: AT2G39270, AT3G54826, AT3G22330, AT1G30680, AT1G72330, AT3G62530, AT4G30920, AT5G17340, AT2G37230, AT2G47510, AT4G23760, AT2G23370, AT4G27585. ii) proteins with operational Cys-based redox switches, which were previously identified in seed proteomes (24). (**B**) Amino acid sequences of the peptides involved in intermolecular disulfide formation of the five Arabidopsis alternative oxidases isoforms. Multiple sequence alignment of the alternative oxidase (AOX) isoforms AOX1a-d & AOX2. The Cys-residues responsible for dimer formation highlighted in red, neighbouring trypsin-cleavage sites (K & R residues) highlighted in blue.

**Table S1. Statistical analysis of germination assays of *Arabidopsis* seeds after controlled deterioration.** *Arabidopsis* seeds were stored for 0, 5, 12, 19 and 26 days at 37 °C and 75 % relative humidity. Seeds were then germinated on plates either in the absence of sucrose (-sucrose) or supplemented with 1 % (w/v) sucrose (+ sucrose). According to two-way ANOVA and Tukey`s multiple comparison test, the *p*-values for comparisons with germination percentages of Col-0 are given: $p > 0.05$ in grey and $p < 0.05$ in black.

**- Sucrose**

| | Days after sowing | | | | | | | | | | | |
|---|---|---|---|---|---|---|---|---|---|---|---|---|
| | Day 2 | | | Day 3 | | | Day 4 | | | Day 5 | | |
| Days of aging | *gr2* | *trx-o1* | *ntr a/b* | *gr2* | *trx-o1* | *ntr a/b* | *gr2* | *trx-o1* | *ntr a/b* | *gr2* | *trx-o1* | *ntr a/b* |
| 0 | 0.7713 | < 0.0001 | 0.1024 | 0.9984 | 0.0526 | 0.9829 | > 0.9999 | 0.9342 | 0.9924 | > 0.9999 | 0.9981 | 0.9951 |
| 5 | 0.0311 | < 0.0001 | 0.3669 | 0.5502 | 0.2596 | > 0.9999 | 0.9497 | 0.7979 | > 0.9999 | 0.9998 | 0.9975 | > 0.9999 |
| 12 | 0.3642 | 0.0015 | 0.0779 | 0.9947 | < 0.0001 | 0.0800 | 0.9754 | < 0.0001 | 0.7345 | 0.9840 | 0.0003 | 0.9949 |
| 19 | > 0.9999 | > 0.9999 | > 0.9999 | < 0.0001 | < 0.0001 | < 0.0001 | < 0.0001 | < 0.0001 | < 0.0001 | < 0.0001 | < 0.0001 | < 0.0001 |
| 26 | > 0.9999 | > 0.9999 | > 0.9999 | > 0.9999 | > 0.9999 | > 0.9999 | > 0.9999 | 0.9992 | > 0.9999 | 0.9996 | > 0.9999 | 0.9996 |

**+ Sucrose**

| | Days after sowing | | | | | | | | | | | |
|---|---|---|---|---|---|---|---|---|---|---|---|---|
| | Day 2 | | | Day 3 | | | Day 4 | | | Day 5 | | |
| Days of aging | *gr2* | *trx-o1* | *ntr a/b* | *gr2* | *trx-o1* | *ntr a/b* | *gr2* | *trx-o1* | *ntr a/b* | *gr2* | *trx-o1* | *ntr a/b* |
| 0 | < 0.0001 | < 0.0001 | > 0.9999 | 0.9984 | 0.6108 | 0.2045 | 0.9998 | 0.8768 | 0.2997 | > 0.9999 | 0.9664 | 0.9415 |
| 5 | 0.0147 | < 0.0001 | 0.9995 | 0.9499 | 0.1210 | 0.7252 | 0.9997 | 0.8999 | 0.9841 | > 0.9999 | > 0.9999 | 0.9997 |
| 12 | 0.2505 | 0.0274 | 0.1871 | < 0.0001 | < 0.0001 | < 0.0001 | 0.1186 | < 0.0001 | 0.1790 | 0.6027 | 0.2409 | 0.8563 |
| 19 | > 0.9999 | > 0.9999 | > 0.9999 | 0.1727 | 0.1727 | 0.8897 | < 0.0001 | < 0.0001 | < 0.0001 | < 0.0001 | < 0.0001 | < 0.0001 |
| 26 | > 0.9999 | > 0.9999 | > 0.9999 | 0.9998 | > 0.9999 | 0.9998 | 0.9997 | > 0.9999 | 0.9997 | 0.9997 | > 0.9999 | 0.9997 |

**Dataset S1 (separate file)**

IodoTMT-based Cys redox proteomic dataset.
